# Mapping mitonuclear epistasis using a novel recombinant yeast population

**DOI:** 10.1101/2022.08.31.505974

**Authors:** Tuc H.M. Nguyen, Austen Tinz-Burdick, Meghan Lenhardt, Margaret Geertz, Franchesca Ramirez, Mark Schwartz, Michael Toledano, Brooke Bonney, Benjamin Gaebler, Weiwei Liu, John F. Wolters, Ken Chiu, Anthony C. Fiumera, Heather L. Fiumera

## Abstract

Natural genetic variation in mitochondrial and nuclear genomes can influence phenotypes by perturbing coadapted mitonuclear interactions. Mitonuclear epistasis, i.e. non-additive phenotype effects of interacting mitochondrial and nuclear alleles, is emerging as a general feature in eukaryotes, yet very few mitonuclear loci have been identified. Here, we present a novel advanced intercrossed population of *S. cerevisiae* yeasts, called the Mitonuclear Recombinant Collection (MNRC), designed explicitly for detecting mitonuclear loci contributing to complex traits, and use this population to map the genetic basis to mtDNA loss. In yeast, spontaneous deletions within mtDNAs lead to the *petite* phenotype that heralded mitochondrial research. We show that in natural populations, rates of *petite* formation are variable and influenced by genetic variation in nuclear, mtDNAs and mitonuclear interactions. We then mapped nuclear and mitonuclear alleles contributing mtDNA stability using the MNRC by integrating mitonuclear epistasis into a genome-wide association model. We found that associated mitonuclear loci play roles in mitotic growth most likely responding to retrograde signals from mitochondria, while associated nuclear loci with main effects are involved in genome replication. We observed a positive correlation between growth rates and *petite* frequencies, suggesting a fitness tradeoff between mitotic growth and mtDNA stability. We also found that mtDNA stability was influenced by a mobile mitochondrial GC-cluster that is expanding in certain populations of yeast and that selection for nuclear alleles that stabilize mtDNA may be rapidly occurring. The MNRC provides a powerful tool for identifying mitonuclear interacting loci that will help us to better understand genotype-phenotype relationships and coevolutionary trajectories.

**Author Summary:** Genetic variation in mitochondrial and nuclear genomes can perturb mitonuclear interactions and lead to phenotypic differences between individuals and populations. Despite their importance to most complex traits, it has been difficult to identify the interacting loci. Here, we created a novel population of yeast designed explicitly for mapping mitonuclear loci contributing to complex traits and used this population to map genes influencing the stability of mitochondrial DNA (mtDNA). We found that mitonuclear interacting loci were involved in mitotic growth while non-interacting loci were involved in genome replication. We also found evidence that selection for mitonuclear loci that stabilize mtDNAs occurs rapidly. This work provides insight into mechanisms underlying maintenance of mtDNAs. The mapping population presented here is an important new resource that will help to understand genotype/phenotype relationships and coevolutionary trajectories.

## Background

Interactions between mitochondrial and nuclear genomes are essential for the mitochondrial functions that power eukaryotic life. Mitonuclear interactions can be direct, as physical contacts between mitochondrial and nuclear genes and their products are needed for mitochondrial DNA (mtDNA) replication and maintenance, transcription, translation, assembly and function of mitochondrially encoded components of oxidative phosphorylation (OXPHOS) complexes [1]. Mitonuclear interactions can also be indirect, though anterograde (nucleus-to-mitochondria) and retrograde (mitochondria-to-nucleus) signaling where metabolites, biochemicals or RNAs direct gene expression in response to metabolic needs or environmental stressors [2, 3]. Genetic variation in natural systems can alter the efficiencies of these interactions leading to phenotypic differences between individuals and populations. Mitonuclear epistasis, defined as the non-additive phenotypic effects of interacting mitochondrial and nuclear allele pairs, has been demonstrated across Eukarya, including vertebrates [4–6], invertebrates [7–12], plants [13] and fungi [14–20]. In humans, allelic variation in mitonuclear interactions contributes to human diseases [21–25]. Identification of important mitonuclear loci will provide insight into evolutionary and coevolutionary processes and facilitate mitochondrial disease prediction and treatments.

Selection for mitonuclear interactions could contribute to speciation [26, 27]. Because of the large interest in uncovering speciation loci, strategies to identify mitonuclear epistasis often focus on analyzing mitonuclear incompatibilities in interspecific and inter-population hybrids or in mitonuclear hybrids where nuclear genomes from one population are paired with the mtDNAs from another. Sometimes candidate genes for these incompatibilities can be revealed through deductive reasoning. For example, in crosses between populations of *Tigriopus californicus* where mtDNA inheritance was controlled, F2 hybrids showed mitonuclear-specific OXPHOS enzyme activities and mtDNA copy number differences, prompting investigations into mitochondrially encoded electron transport proteins and the mtRNA polymerase [28, 29]. In *Drosophila*, a mtDNA from *D. simulans* paired with a nuclear genome from *D. melanogaster* resulted in mitonuclear genotype with impaired development and reproductive fitness [30]. Fortuitously for mapping purposes, the mtDNA sequences in this mitonuclear panel had very few polymorphisms, enabling the causative alleles behind this incompatibility (a mitochondrially encoded tRNA and a nuclear encoded tRNA synthetase) to be identified [31].

Genetic approaches can also be used to reveal mitonuclear epistatic loci. Chromosomal replacements in interspecific *Saccharomyces* yeast hybrids, followed by plasmid library screening, allowed the identification of mitonuclear incompatibilities between nuclear genes encoding intron splicing factors from one species and their mitochondrially-encoded targets in the other [17, 32, 33]. Quantitative trait loci (QTL) mapping approaches using genotype/phenotype associations of recombinant progeny containing different mtDNAs have also been used to seek intraspecific mitonuclear incompatibilities [34–37]. Analysis of meiotic segregants following forced polyploidy enabled the detection of QTLs for interspecific mitonuclear incompatibilities contributing to sterility barriers in *Saccharomyces* [38]. Due to the low resolution of QTL mapping, specific loci were not identified, but in some cases, regions of mitonuclear genomic interest were implicated. Other approaches to uncovering mitonuclear epistatic loci include differential expression analysis [5, 39–41] and experimental evolution [42, 43]. Because relatively few genetic backgrounds are used for these mapping approaches, even if single gene pairs are identified, it is not clear if these approaches will reveal a general picture of mitonuclear epistasis or lineage specific idiosyncrasies.

Mitonuclear epistasis could explain over 30% of the phenotypic variances observed in a panel of *S. cerevisiae* yeasts consisting of 225 unique mitonuclear genotypes [15]. In the current study, our goal was to map naturally occurring alleles leading to mitonuclear epistasis in this yeast population. Detecting mitonuclear epistatic loci (or any g × g interaction) using association approaches is challenging due to low allele frequencies, the large number of tests required to detect pairwise epistasis and the potential for the environment to affect genetic interactions [44–46]. To overcome some of these challenges, we created a recombinant population of *S. cerevisiae* designed explicitly to detect mitonuclear epistasis through association testing. We used controlled and random matings with 25 natural yeast isolates to create a multiparent advanced intercross population in order to reduce effects of the known population structure in yeast [47] and allow for finer mapping resolution. We controlled the mtDNA inheritance in the recombinant collection such that strains shared a single mitotype, and then replaced this mitotype with two different mtDNAs resulting in a collection of nuclear genotypes paired with three different mitotypes. This should increase power to detect mitonuclear interactions across the full mapping population and to detect nuclear effects within a given mitotype. Mitonuclear interactions could thus be integrated into phenotype/genotype association models to detect loci contributing to complex traits.

We focused on mapping alleles contributing the stability of the mitochondrial genome, a complex trait controlled by many nuclear loci [48–52]. Large scale deletions within mtDNAs are a common form of mtDNA instability, leading to mitochondrial heteroplasmies (defined as having more than one mitotype present) and deterioration of organismal health [53–57]. *Saccharomyces* yeasts do not tolerate mtDNA heteroplasmies and will fix for a single mitotype after just a few mitotic divisions [58]. Yeast cells containing these spontaneous mtDNA deletions can grow via fermentation and will form small, *petite*, colonies in comparison to the larger *grande* colonies formed by respiring cells, making it possible to quantify rates of mtDNA deletions [59]. Laboratory strains have accumulated multiple genetic variants leading to increased *petite* frequencies [51]. We reasoned that natural genetic variation would lead to differences in *petite* frequencies among wild isolates of *S. cerevisiae* through mitonuclear epistasis and that our recombinant population would enable the identification mitonuclear epistatic loci.

Here, we show that mtDNA stability in *S. cerevisiae* populations is influenced by mitonuclear epistasis and by independent effects of both nuclear and mitochondrial genomes. We describe the construction of the Mitonuclear Recombinant Collection, and how we were able to use this to detect nuclear loci that participate in the mtDNA stability through main (independent) effects and mitonuclear interactions. The epistatic loci identified revealed that growth rates influence the stability of mtDNAs. The mitonuclear recombinant collection provides a new tool for identifying mitonuclear loci that are important in nature.

## Results

### mtDNA stability is a quantitative trait influenced by mitochondrial GC clusters and mitonuclear interactions

Given the mechanistic complexities of mtDNA replication and maintenance, we hypothesized that standing genetic variation would contribute to differences in mtDNA stability. To test this, we examined the frequencies of spontaneous *petite* colony formation in 21 isolates of *S. cerevisiae* (**Figure 1**, **Table S1**). Rates of mtDNA deletions in these isolates ranged from 0.3 to 9.9%. All 21 isolates had lower *petite* frequencies than S288c, a widely used laboratory strain known to harbor nuclear variants that promote exceptionally high rates of *petite* formations [51]. Within the wild isolates, genetic variation within and between broadly-defined ancestral populations contributed to differences in the rates of mtDNA loss (*P*<0.001 for strain and population, **Table S2**). Strains with Wine/European or mosaic origins had the lowest rates of *petite* formation, while strains with West African or Sake/Asian origins had the highest.

**Figure 1.**
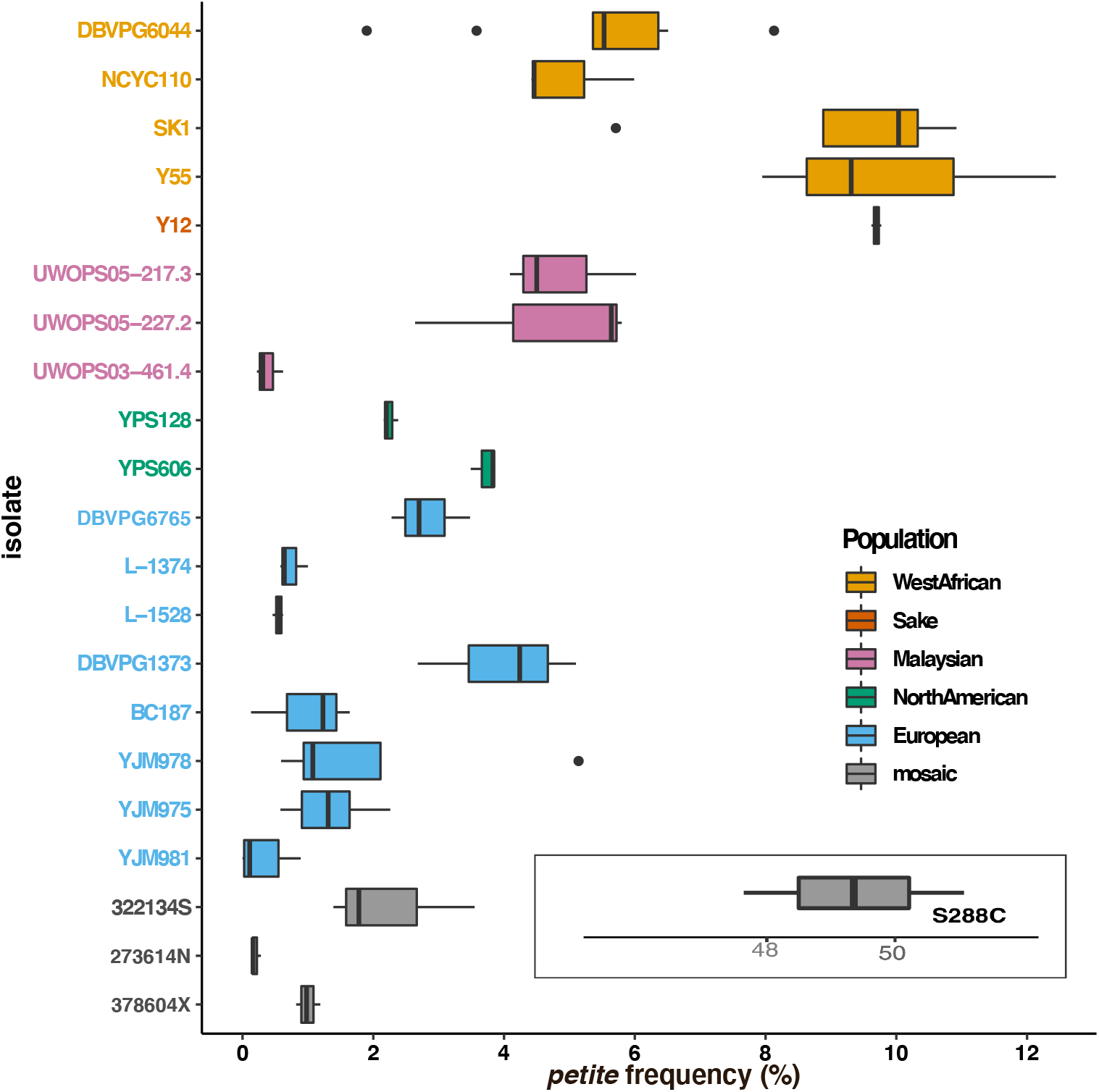
Genetic variation contributes to *petite* frequency. Boxplots showing *petite* frequencies (total *petite* colonies/total CFU *100) are presented for haploid derivatives of 21 wild *S. cerevisiae* isolates and the reference strain, S288c. Strains are colored by broad population identities from [47, 60, 61] as indicated in the key. The *petite* frequency for the reference strain is offset for scaling purposes. *Petite* frequencies differ between populations and between strains within populations (ANOVA, **Table S2**).

The differences in *petite* formation rate between wild isolates could be due to genetic variation in mtDNAs, nuclear genomes and/or mitonuclear interactions. To examine the independent effects of different mitotypes on mtDNA stability, we introduced 18 different mtDNAs into a common nuclear background and assayed rates of mtDNA deletions. In these iso-nuclear strains, *petite* frequencies ranged from 5.5 to 31.0 due to differences in mitotypes (ANOVA, *P*<0.001, **Table S3**). To determine if *petite* formation correlated with any particular mitochondrial feature, we analyzed the mtDNAs for their length, intron content, GC% and GC-cluster content (**Table S4**). The GC-rich regions of yeast mtDNAs are predominantly due to short (~30-40 bp) mobile GC-rich palindromic sequences called GC-clusters that interrupt long intergenic AT-rich sequences [62, 63]. GC content and GC cluster numbers were tightly correlated (r = 0.95, *P*<0.001), and both were correlated with the total length of mtDNAs (r = 0.59, *P*=0.001 and r=0.70, *P*<0.001, respectively). *Petite* frequencies, however, did not correlate with the total length of mtDNAs (r=0.12, *P*=0.63) nor the total lengths of intron sequences (*r=−0.09*, *P*=0.72). Thus, it is likely that GC rich regions play an important role in mtDNA stability. In fact, we found that *petite* frequencies correlated with overall GC% of mtDNAs (Pearson’s r=0.59, *P*=0.013, **Fig 2A**) but not the total numbers of GC clusters (r=0.44, *P*=0.07), mtDNA length (r=0.12, *P*=0.63) nor total length of intron sequences (r=−0.09, *P*=0.72). A presumed mechanism for *petite* formation is illegitimate recombination in mitochondrial GC clusters [59, 64, 65]. To see if any particular type of GC cluster could explain mtDNA instabilities, we sorted the GC clusters into 9 classes described by sequence homologies [66]. We found that the M4 GC-cluster class, but no other, correlated with *petite* frequencies (r=0.67, *P*=0.004, **Fig 2B**). This particular cluster appears to be expanding in the mtDNAs of strains with West African lineages [67] and explain the observed correlation.

**Figure 2.**
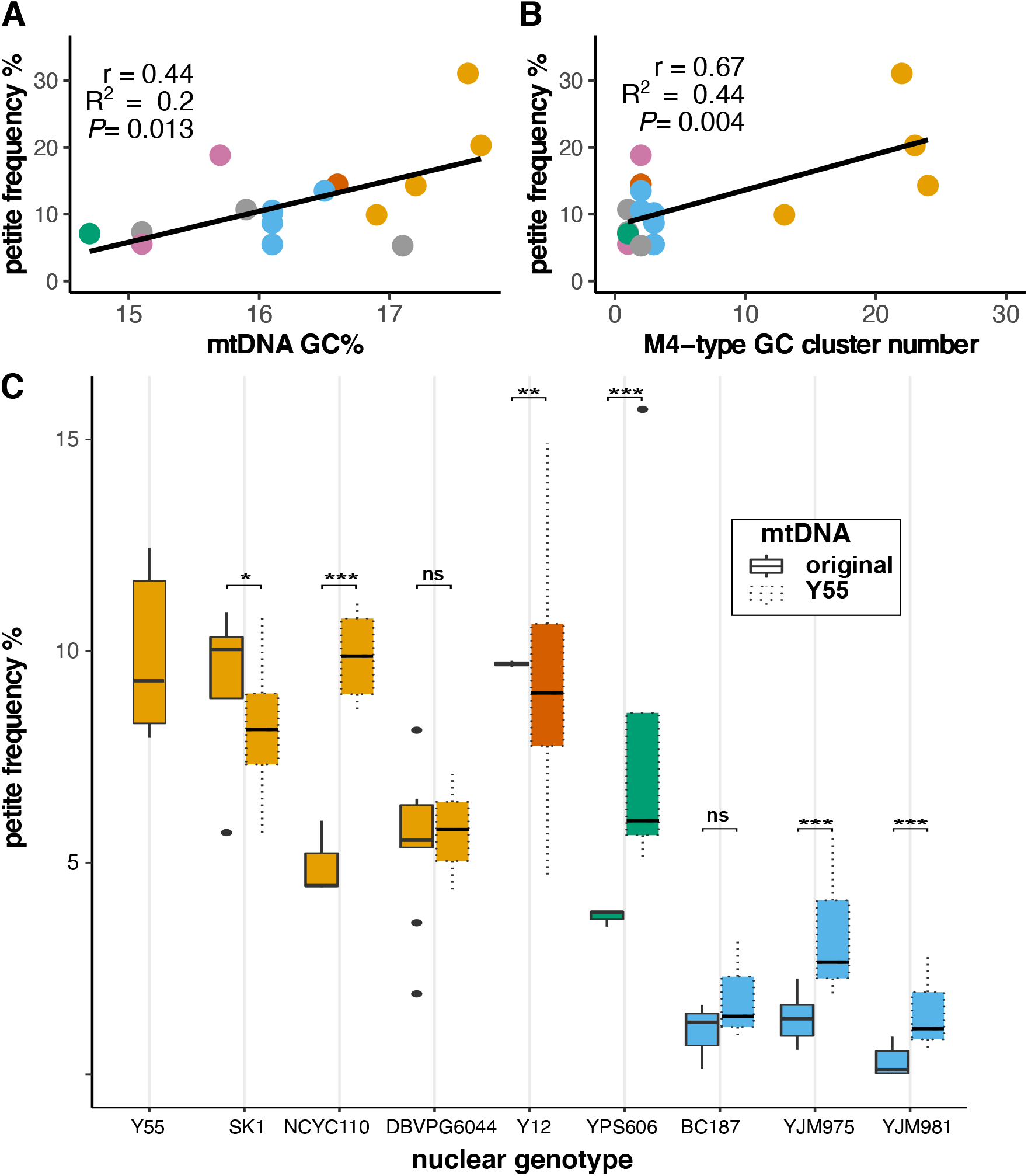
Mitochondrial GC-content influences stability of mtDNAs. *Petite* frequencies of iso-nuclear strains (containing the nuclear genome from strain Y55) correlate with **A**. total GC% of mtDNAs and **B**. the numbers of M4-type GC clusters. Pearson’s correlation coefficient (r), coefficient of determination (R^2^) and *P-*value significance are shown. **C**. *Petite* frequencies for strains containing original (solid outlines) or the GC-cluster rich mtDNA from Y55 (dotted outlines) are shown as box and whisker plots. Colors indicate nuclear genotype population as described in **Fig 1**. Significance of individual ANOVAs comparing the original and synthetic mitonuclear genotypes are shown. * *P<0.05, \**\* *P* ≤ 0.005, *** *P* ≤ 0.001.

GC clusters, however, do not solely control rates of *petite* formation. When a mtDNA from the West African strain with the highest *petite* frequency was introduced into different nuclear backgrounds, *petite* frequencies were increased in 4 of 8 nuclear backgrounds tested and decreased in 2 of them (**Fig 2C**). This is consistent with both nuclear and mitonuclear effects. Nuclear backgrounds from Wine/European origins maintained relatively low *petite* frequencies even when harboring this GC-cluster rich mtDNA, indicating that nuclear genotypes largely control mtDNA stability, at least for these strains.

Some of the synthetic mitonuclear combinations led to even higher *petite* frequencies than observed in the original isolates suggesting that mitonuclear interactions also play a role in mtDNA stability. To formally test for mitonuclear effects on mtDNA stability, we examined *petite* frequencies in 16 mitonuclear cybrids created by exchanging mtDNAs between 4 strains with West African lineages (4 nuclear × 4 mtDNAs). A two-way ANOVA showed that mtDNA stability was influenced by nuclear genotypes, mitotypes, and mitonuclear interactions (*P*<0.001 for each term, **Table S5**), where certain mitonuclear combinations showed very large increases in the rates of *petite* formation (**Fig 3**). Within these strains, the original mitonuclear genotypes had lower *petite* frequencies than any of their synthetic mitonuclear combination (**Fig S1**). Given the recent expansion of the destabilizing M4 clusters within the West African strains, this observation suggests that selection for mitonuclear interactions that stabilize mtDNAs has occurred quickly in ways that are strain-specific.

**Fig 3.**
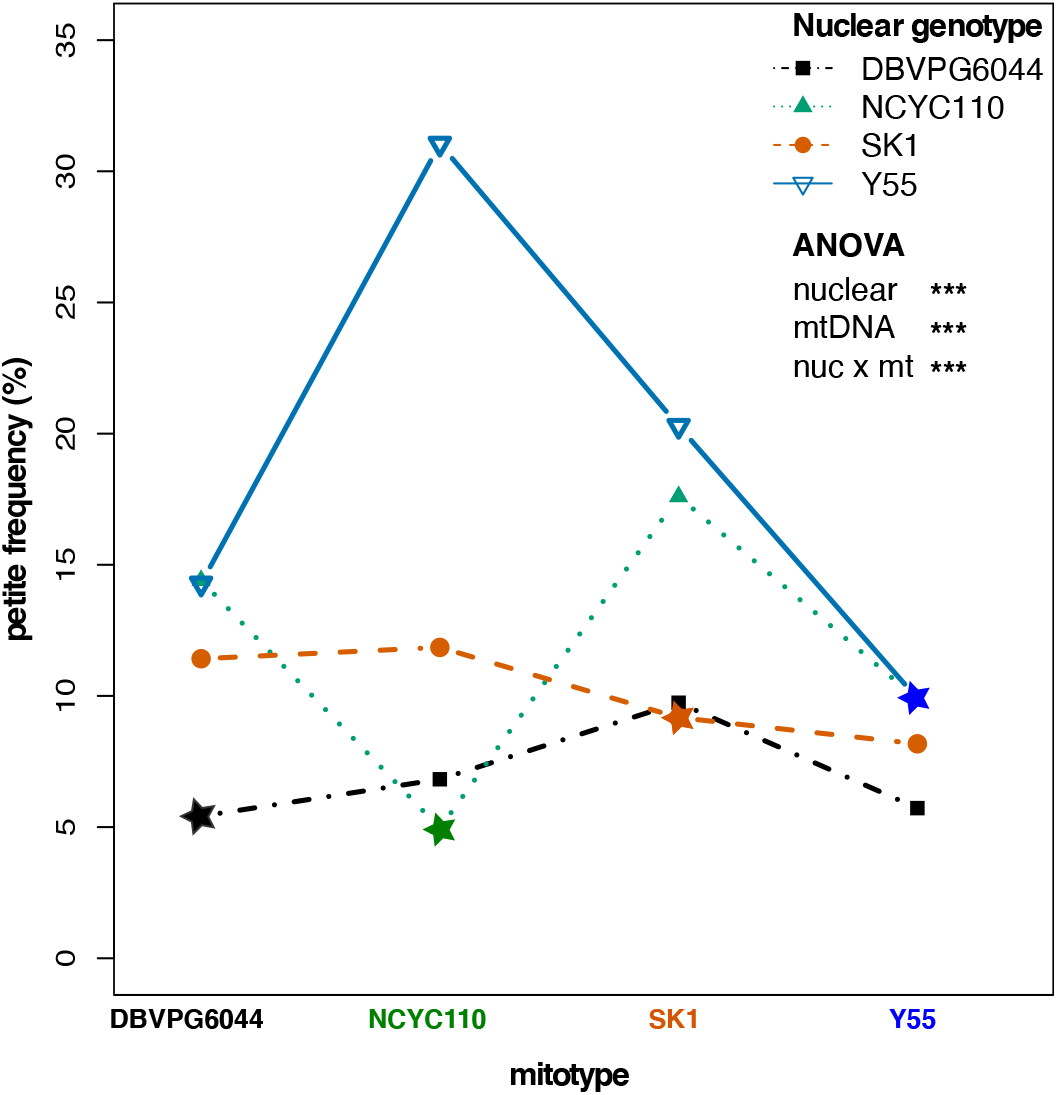
Coadapted mitonuclear interactions stabilize mtDNAs. *Petite* frequencies for strains containing 4 nuclear and 4 mtDNA backgrounds are shown as an interaction plot. Colored lines connect a nuclear genotype paired with different mitotypes. The original mitonuclear genotype combination is starred. The nonparallel lines indicate mitonuclear epistasis. ANOVA revealed highly significant nuclear, mtDNA, and mitonuclear genotypes contributions (**Table S5**). *** *P* ≤ 0.001.

### Creation of a Mitonuclear Recombinant Collection for association studies

Mitonuclear interactions explain a significant proportion of phenotypic variances in *S. cerevisiae* yeasts and involve numerous, as yet unmapped, loci [14, 15, 68]. We sought a general genome-wide mapping approach that would facilitate the mapping of both the nuclear and mitonuclear loci underlying complex traits such as mtDNA stability. We created a multiparent recombinant collection of *S. cerevisiae* strains specifically designed for association mapping of nuclear and mitonuclear loci (called the Mitonuclear Recombinant Collection, or MNRC) (**Fig 4**). To do this, we first replaced the mtDNAs in 25 wild divergent yeast isolates such that each contained an identical mitotype, and then mated them to create each possible heterozygous diploid. The diploids were sporulated and ~10,000 haploid F1 haploid progeny were isolated and then randomly mated. F1 diploids were then isolated and sporulated. The process was repeated for a total of 7 rounds of meiosis. A collection of 181 F7 haploids (named Recombinant Collection 1 (RC1)) were isolated and fully sequenced. The mtDNAs from RC1 strains were removed (creating RC*ρ*^0^) and replaced with two additional mtDNAs via karyogamy-deficient matings, creating populations RC2 and RC3. The GC cluster content of the mtDNAs in the MNRC are classified as low (117 clusters in RC2), medium (137 in RC1), or high (203 in RC3).

**Figure 4.**
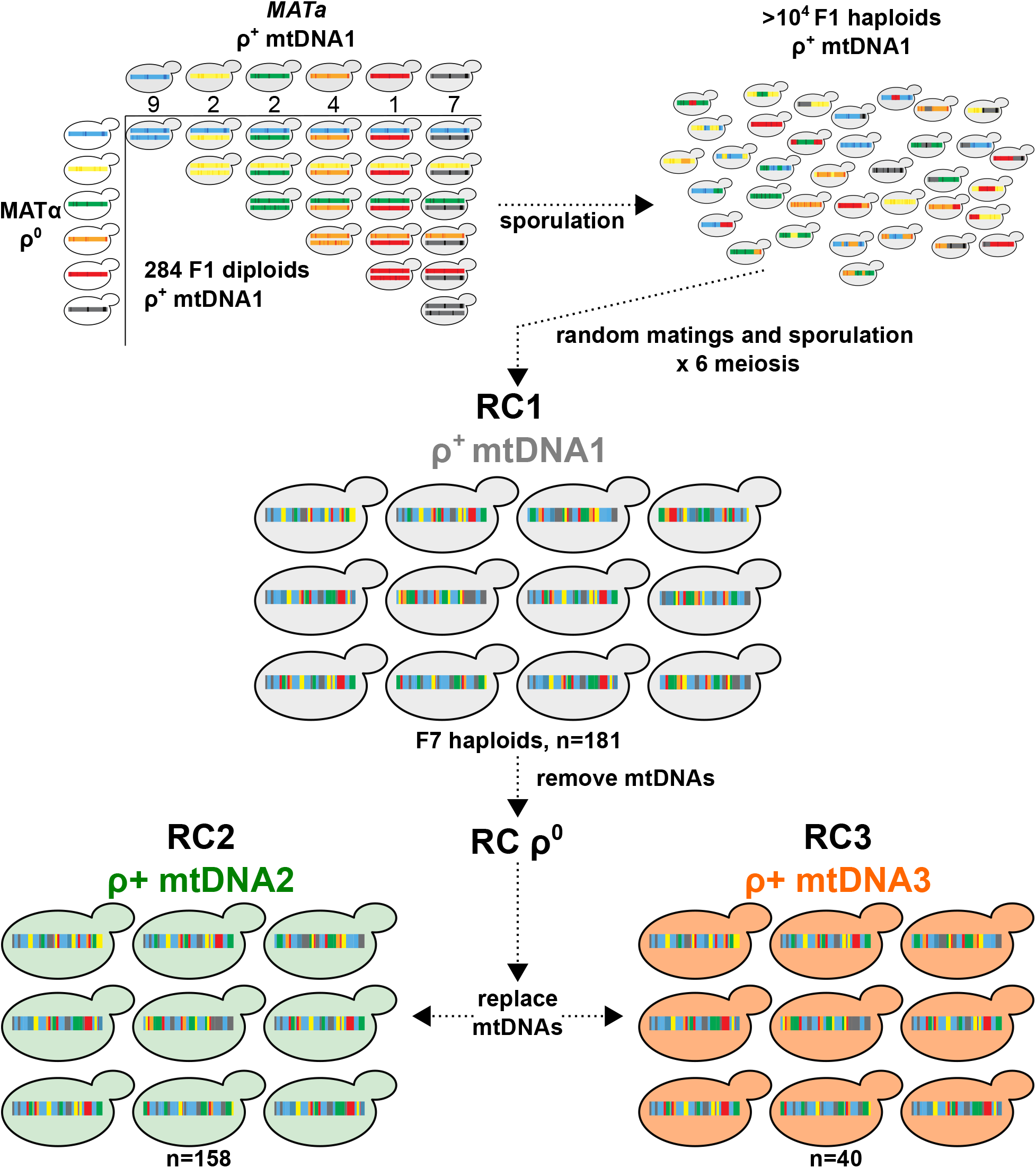
A Mitonuclear Recombinant Collection designed for association mapping. 25 unique genetic backgrounds of *S. cerevisiae* fixed for a single mitotype were systematically crossed to create each possible F1 heterozygous diploid with identical mtDNAs (284 total). The numbers of unique genotypes from each parental ancestral background are indicated; blue (Wine/European); yellow (Malaysian); green (North American); orange (West African); black (mosaic). The F1 diploids were sporulated and F1 recombinant haploid progeny isolated. Following 6 rounds of random matings, haploid F7 progeny were isolated to create RC1. The mtDNAs from RC1 were removed (RC1*ρ*^0^) and replaced with 2 different mtDNAs, creating RC2 and RC3. The mtDNAs is RC1,2, and 3 are from the wild isolates 273614N, YPS606, and NCYC110, respectively.

To generate SNPs tables for association testing, RC1 strains were sequenced to ~40x coverage. Paired-end reads were aligned to the *S. cerevisiae* reference genome and the locations of nuclear SNPs and small indels were extracted from each alignment. Polymorphic sites were filtered by removing telomeric regions and SNPs/indels with low allele frequencies (MAF <5%). Following filtering, 24,955 biallelic sites across the 16 yeast chromosomes with an average of ~2200 SNPs/chromosome were available for association testing. Chromosomal polymorphic data are summarized in **Table S6**. Our read alignments and subsequent analyses did not account for chromosomal rearrangements, such as copy number variants, translocations and inversions, that would map to similar locations of the reference genome nor genomic regions absent in the reference strain.

We validated that this novel recombinant population could be used for simple association studies. Strains from RC1 were phenotyped for growth on copper sulfate and an association test was performed to identify SNP variants associated with copper tolerance. A single peak on Chr. 8 coincided with a region containing *CUP1*, the copper binding metallothionein (**Fig S2**). Variation at this locus is known to lead to high copper tolerances found in Wine/European isolates [69] and has previously been identified through association studies using wild isolates [47, 70]. Thus, the recombinant collection produced here is successful for association studies despite a relatively low number of parental strains.

### Nuclear and mitonuclear associations for mtDNA stability

To map nuclear and mitonuclear associations, *petite* frequencies were collected for each strain in RC1, RC2 and RC3. In RC1, the *petite* frequencies ranged from 0.0% to 27.7% forming a continuum, as would be expected for a complex trait involving numerous loci (**Fig 5A**). The same rank orderings were not observed in RC2 or RC3, revealing the influences of mitonuclear interactions. RC3, containing the GC cluster rich mtDNA, had a higher average *petite* frequency than RC1 or RC2 (**Fig S3**).

**Figure 5.**
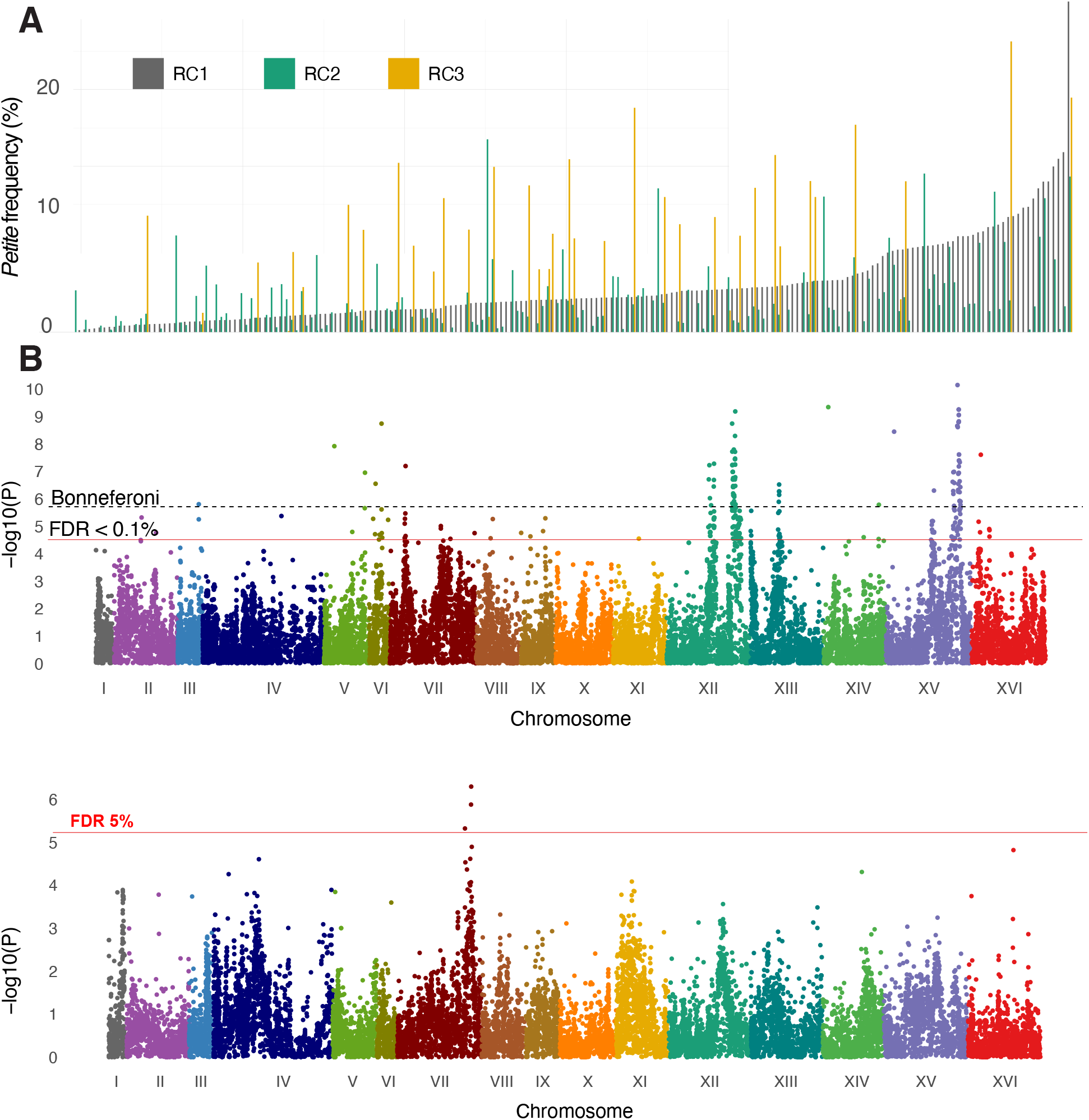
Nuclear SNPs associate with mtDNA stability through main effects and mitonuclear interactions. **A.** *Petite* frequency is a quantitative trait influenced by natural genetic variation. *Petite* frequencies for strains from RC1 (gray) were rank ordered from lowest to highest. The *petite* frequencies for strains from RC2 (green) and RC3 (gold) are plotted next to isonuclear strains from RC1. **B**. Manhattan plots show main effect nuclear SNPs associated with *petite* frequencies in ways that were independent of mtDNA and **C**. nuclear SNPs whose association is dependent on mitotype (ie. mitonuclear). The different plot profiles indicate that the main effect nuclear SNPs are different that those involved with mitonuclear interactions. FDR thresholds at 0.1% (*P* < 4.1 × 10^−5^) for nuclear associations and 5.0% (*P* < 1.2 ×10^−5^) for mitonuclear associations and a conservative Bonferroni threshold (*P* < 2.0 × 10^−6^) are shown.

In theory, the recombinant genomes and fixed mitotypes of the RC strains should reduce effects of population structure, improve statistical power while using a smaller number of samples, limit false positives, and control for mitonuclear interactions. We performed association testing to identify nuclear loci that had both a main effect and interacted with the mtDNA to influence mtDNA stability. Mating types, auxotrophic markers and residual population structure as determined by principal component analyses were included as covariates (see METHODS). The significance profiles of associations for nuclear variants that were independent (*nuclear SNP*, **Fig 5B**) and dependent on mitotype (*nuclear SNP× mtDNA*, **Fig 5C**) were different, providing confidence that the association model is able to detect nuclear features that are unique to either main or epistatic effects.

Nuclear SNPs whose effects were independent of mitotype resulted in stronger associations than mtDNA-dependent alleles. This is not surprising given that independent contributions of nuclear genotypes influence growth phenotypes to a greater extent than mitonuclear interactions [14, 15, 68]. At a false discovery rate (FDR) < 0.1% (Q-value = 0.001), we observed 130 mtDNA-independent associated SNPs located within or 250 bp upstream of coding sequences (**Table S7**). In comparison, at FDR <5% (Q-value = 0.05), there were 3 mtDNA-dependent associated SNPs (**Table S8**). Strong nuclear effects could mask mitonuclear interactions. We identified the alleles with the strongest effects by calculating the effect size of each SNP with mitotype-independent associations as the difference between the average *petite* frequencies of each allele weighted by its frequency in the recombinant collections (**Table S9**). This revealed that the highest effect size was attributed to a SNP on Chr. 15 predicting a G50D missense mutation in *MIP1*, the mitochondrial DNA polymerase required for replication and maintenance of mtDNA. To improve power of detecting mitonuclear associations, we repeated the analysis including the *MIP1* SNPs as covariates. This removed the mitotype-independent associations on Chr. 15 and increased the numbers of significant mitonuclear associations from 3 to 27 without changing the overall association profiles (**Fig S4, Table S8**).

### Mitonuclear associations: mitotic growth signaling pathways

The 27 mitonuclear SNP associations corresponded to 21 unique genes, including 11 in a QTL on Chr. 7 (**Table S8**). Within this QTL, the three strongest associations corresponded to SNPs near the coding start sites of the genes *HGH1* and *SMI1* and an in-frame deletion within *BNS1*. Interestingly, mitochondrial activities have not been shown for these genes. A second QTL on Chr. 1 contained a SNP upstream of *YAT1*. To verify the involvement of these genes in mitonuclear interactions affecting mtDNA stability, we first looked at how removing each gene influenced *petite* frequencies. In the parental background of the *S. cerevisiae* knockout collection (BY4741) and with the BY4741 mitotype, the null mutant *hgh1*Δ lowered rates of *petite* formation while null mutants *bns1*Δ, *smi1*Δ and *yat1*Δ showed no significant effect (**Fig 6A**). We next introduced two different mitotypes (the GC cluster-rich mitotype NCYC110 used in RC3 and the GC cluster-poor mitotype YPS606 used in RC2) into the parental and deletion strains and measured *petite* frequencies. Mitonuclear interactions were readily observed; the GC-rich mitotype led to higher *petite* frequencies in the *hgh1*Δ and *yat1*Δ deletion strains, whereas the same mitotype led to a lower *petite* frequency in the parental background (**Fig 6B**). Two-way ANOVAs showed significant mitonuclear interactions when comparing the parental strain to each deletion strain with each mtDNA comparison (**Table S10**). Similar mitonuclear effects were also observed for *bns1*Δ and *smi1*Δ when compared to the parental background (**Fig S5, Table S11**). While the high *petite* frequency of the parental strain complicates the interpretation of these assays, the differential responses provide strong evidence that the association model was successful in identifying genes that influence mtDNA stability through mitonuclear epistatic interactions.

**Figure 6.**
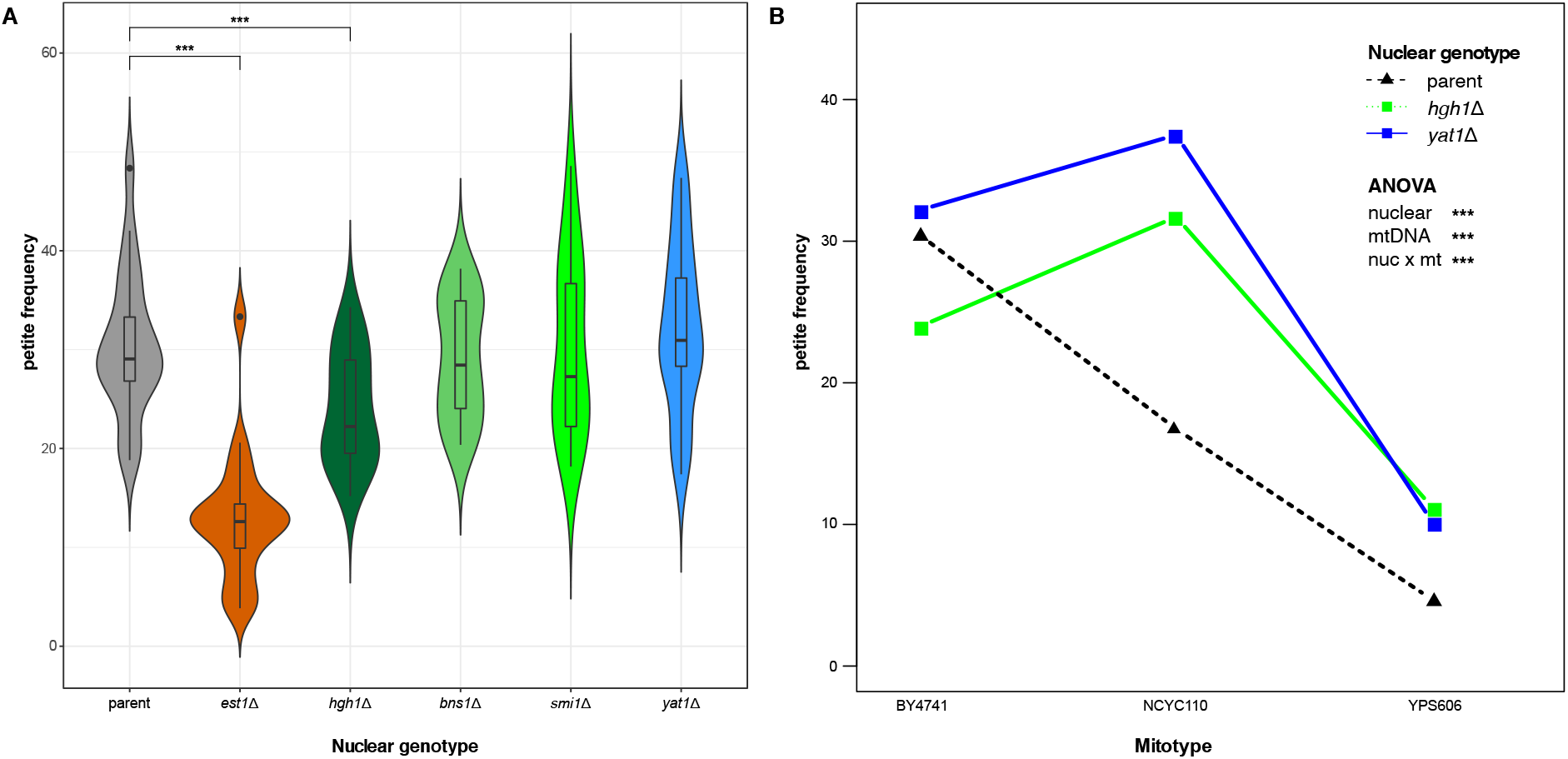
Mitonuclear interactions influencing mtDNA stability include *HGH1* and *YAT1*. **A.** *Petite* frequencies for strains lacking genes with mitotype-independent (*est1Δ)* and mitonuclear associations (*hgh1*Δ, *bns1*Δ, *smi1*Δ and *yat1*Δ) are shown as boxes in violin plots. Significant differences between the parental and each deletion strain are shown. * *P<0.05, \**\* *P* ≤ 0.005, *** *P* ≤ 0.001. **B**. *Petite* frequencies of *hgh1*Δ and *yat1*Δ strains depend on mitotype. Colored lines connect each nuclear genotype paired with different mtDNAs. Non-parallel lines indicate that mitotypes influence mtDNA stability through mitonuclear interactions. Comparisons of the parental and each deletion strain with any 2 different mtDNAs showed highly significant mitonuclear interactions (*P*<0.001 for each comparison, **Table S10**). Mitotype BY4741 is the parental mtDNA in the yeast deletion collection. Mitotypes NCYC110 and YPS606 were used in RC3 and RC2, respectively. A minimum of 20 replicates for all *petite* assays were performed.

We then tested if the mitonuclear associations could be explained by differences in gene expression. mRNAs were extracted from a subset of 18 strains from the RC2 and RC3 collections containing GC-cluster poor and rich mitotypes, respectively. We attempted to control for *MIP1* alleles and the 2 most common SNP haplotypes for mitonuclear candidate genes on Chr. 12 (containing *HGH1, BNS1, and SMI1)* and Chr. 1 (containing *YAT1*). Across the subset of RC2 and RC3 strains containing alternate mitonuclear haplotypes with each *MIP1* allele, *YAT1* SNPs (Chr. 1) were in complete LD with the associated SNPs on Chr. 12 and are thus considered together. Expression of *HGH1* showed a modest positive correlation with *petite* frequencies (r=0.43, *P*=0.08) (**Fig S6**). Expression of *MIP1, YAT1*, *BNS1* and *SMI1* showed slight but non-significant positive correlations (**Fig S6**). In these strains, mitotypes did not influence expression of the mitonuclear candidate genes nor on the expression of the mtDNA polymerase, *MIP1* (**Table S12**). The expression of *MIP1* was, however, influenced by the haplotype of the Chr. 12/1 QTLs (**Fig S7A** while expression of *BNS1* was influenced by *MIP1* alleles (**Fig S7B**, **Table S12**). *BNS1* and *YAT1* expression showed *MIP1* haplotype × mitonuclear haplotype effects. Interestingly, the higher expression in these genes were observed in strains containing the *MIP1* alleles that led to a high *petite* frequency. These data suggest that the genotypes of *MIP1* and the mitonuclear candidate genes may influence the expression of each other to affect mtDNA stability.

The mitonuclear candidate genes, *YAT1, HGH1*, *BNS1*, and *SMI1* are broadly connected to mitotic growth. To investigate a potential relationship between mitotic growth and mtDNA stability, we compared the growth rates of wild yeast isolates and their *petite* frequencies and found that the faster growing strains were more likely to lose their mtDNAs (**Fig 7A**). Increasing growth rates by changing glucose availability or temperature also led to an increase in *petite* frequencies (**Fig 7B)**. This suggests a fitness tradeoff between rapid cell growth and deletions of mtDNA. We also noted that increase in *petite* frequencies of the West African strains with synthetic mitonuclear genotypes (**Fig S1**) coincided with an increase in growth rates (**Fig S8**). Selection for mitonuclear interactions that stabilize mtDNAs may act by lowering overall growth.

**Figure 7.**
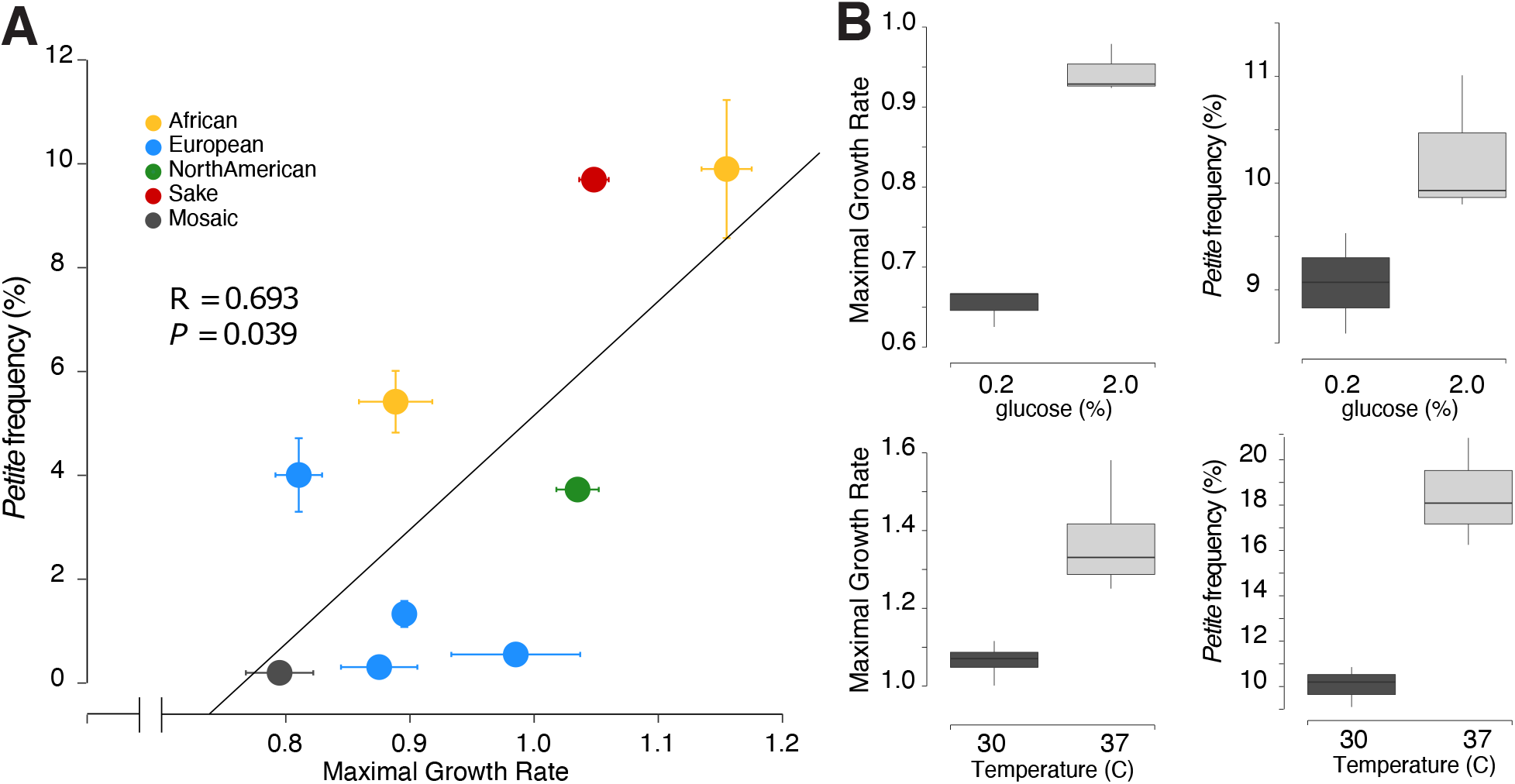
Rapid mitotic growth increases rates of mtDNA loss. **A.** Growth rates of wild isolates and spontaneous *petite* formation are positively correlated. Maximal growth rates (from [14]) of strains grown in conditions used for *petite* assays were plotted against rates of spontaneous *petite* formation. **B**. Growth rates and *petite* frequencies increased as a result of increased glucose concentration or temperature. Assays were performed on Sake strain Y12 (NCYC3605) a minimum of three times. Growth rates are reported as mOD/min.

### Mitotype-independent associations: mtDNA polymerase and a G-quadruplex stabilizing protein

In addition to identifying mitonuclear loci, our model identified 301 SNP whose associations were independent of mitotype. Of these, 130 SNPs were located within 250 bp upstream of coding sequences for 86 unique genes (**Table S7)**. Gene Ontology (GO) analyses revealed enrichments in mitochondrially-targeted genes (27 genes), genes involved in mitochondrial organization (9 genes) and homeostatic processes (10 genes) (**Table S13**). Sixteen of the associated genes generate non-respiratory phenotypes when deleted from the genome [49, 71–73]. In all, 33 (38.4%) of the mitotype-independent associations have known mitochondrial activities.

A large number of the associated SNPs occurred within QTLs on chromosomes 12, 13 and 15 (**Fig 5B**) and were likely associated through linkage. To help identify causative loci, effect sizes for each SNP were calculated as the difference between the average *petite* frequencies of each allele weighted by its frequency in the recombinant collections (**Table S9**). The highest effect size was attributed to a SNP on Chr. 15 predicting a G50D missense mutation in *MIP1*, the mitochondrial DNA polymerase required for replication and maintenance of mtDNA. Associations for two additional *MIP1* missense variants (T540M and H541N) were also significant. High effect sizes were also seen in associated SNPs in genes flanking *MIP1*, including *FRT1* and *PDR10*, and essential genes *KRE5* and *ALA1*. Deletions of *FRT1* or *PDR10* did not significantly alter *petite* frequencies suggesting that these associations are the result of linkage to *MIP1* (**Fig S9**).

Thirty-two associated genes were located within two QTLs on Chr. 12. Here, the largest effect size was for a putative missense variant (L13I) in *EST1* (**Table S9**), encoding a telomerase accessory protein that helps to stabilize of G-quadruplex structures in telomeres during certain stages of the cell cycle [74]. We found that a deletion of *EST1* significantly reduced *petite* frequencies as compared to the parental strain in the yeast knockout collection (**Fig 6A**) while a deletion of *TOP3*, a candidate gene tightly linked to *EST1*, did not (**Fig S9**). This suggests that *EST1* variants likely drive the associations in this QTL. A second QTL on Chr. 12 contained associations for 5 genes with known mitochondrial functions. *ILV5* is a nonspecific mtDNA binding protein involved in the packaging of mtDNA into nucleoids and is required for mtDNA stability [75]. *NAM2*, a mitochondrial tRNA synthetase [76], and *SSQ1*, a mitochondrial chaperone [77], are required for growth via mitochondrial respiration. We found that a deletion of *MDM30*, a component of the ubiquitin ligase complex involved in mitochondrial morphology and turnover [72], lowered *petite* frequencies while a deletion of *ATG33*, a protein involved in mitochondrial mitophagy (Kanki *et al*. 2009), had no effect (**Fig S9**). A single QTL on Chr. 13 contained associations for 6 genes, four of which encode mitochondrially targeted proteins. *COX14* and *SAM37*, and *SOV1* are required for mitochondrial respiration [49, 71, 72] and *RNA14* is essential for growth. Deletions of a 5^th^ gene, *CSM3*, did not significantly alter *petite* frequencies (**Fig S9**).

## Discussion

### Mitonuclear Recombinant Collection

We created a multiparent advanced intercross population (Mitonuclear Recombinant Collection or MNRC) to identify mitonuclear interactions driven by allelic variation in *S. cerevisiae* yeasts through association testing. The MNRC incorporates the genetic variation from 25 wild isolates into ~200 fully sequenced recombinant strains. These strains were paired with three different mitotypes, enabling the detection of mitonuclear interactions via genome-wide association tests. Using the MNRC, we were able to identify loci contributing to mtDNA stability through independent (main) effects and mitonuclear interactions.

The MNRC was designed to help overcome many of the challenges of detecting epistatic interactions. Multiple rounds of random mating should lower existing LD created by the strong population structure in *S. cerevisiae* [47, 61] and reduce the numbers of strains needed for association testing. The MNRC strains are haploid, eliminating dominance effects. Because novel mtDNAs created through mitochondrial recombination may produce unexplained phenotypic variances [19, 68], mtDNA inheritance was controlled. By specifically pairing each mitotype with the nuclear recombinants, the power to detect specific mitonuclear loci was increased. Environments can easily be controlled in yeast, eliminating unexplained variances due to mtDNA × nuclear × environment interactions. We showed that the MNRC can successfully detect nuclear alleles that contribute to phenotypes through independent (main) or epistatic (mitonuclear) effects.

### Loci affecting mtDNA stability

#### Mitonuclear interactions

Our model found epistatic associations for SNPs in four genes (*HGH1, SMI1, BNS1*, and *YAT1*) that are broadly related to mitotic growth. We confirmed these associations by showing that mitotypes influence *petite* frequencies differently in null mutations of each gene as compared to a wild type genetic background. *YAT1* encodes a mitochondrial outer membrane protein that participates in mitochondrial metabolism by importing the acyl groups that enter the Krebs cycle for energy production [78, 79]. Industrial yeast strains have likely adapted to high levels of toxic aldehydes produced during ethanol production by increasing *YAT1* copy numbers [80]. The Yat1 protein is phosphorylated [81], suggesting that its activity is regulated through cell signaling. *HGH1* encodes a translation factor chaperone involved in protein synthesis that is activated in response to DNA damage [82] [83]. The molecular function of *SMI1* is unknown, but it is thought to be a member of the cell signaling pathway that regulates cell wall biosynthesis during mitotic growth [84]. Null mutations of *SMI1* have reduced respiratory growth [85], suggesting its activity influences mitochondrial function. During mitosis, *BNS1* participates in the signaling network directing the exit from anaphase [86]. A high-quality mitochondrial proteomics screen found the Bns1 protein in the mitochondrial matrix [87]. Possibly, these genes are involved in mitonuclear epistasis by changing growth parameters in response to retrograde signals produced by mitochondria with damaged mtDNAs. We expect this to be a general trend. Dominant mutations that suppressed the low growth phenotype of strains with mitochondrial ribosomal protein defects also led to higher *petite* frequencies [88].

Mitotic growth also linked mitonuclear epistasis and mtDNA stability when exchanging mtDNAs between strains from West African lineages. This strain grouping had higher *petite* frequencies (**Fig 1**) and growth rates (**Fig 7**) than most other strains. When mtDNAs were exchanged between West African isolates, strains with original, coadapted mitonuclear genome combinations had lower *petite* frequencies (**Fig 3**) and mitotic growth rates (**Fig S8**) than strains harboring introduced, non-coadapted, mtDNAs. This suggests that selection for mtDNA-stabilizing mitonuclear alleles is rapid and may come at the cost of lowering overall growth rates and may even dictate an upper limit for optimal growth. It is interesting to note that mtDNA instability is a hallmark of fast growing human cancer cells [89]. Experimental evolutions aimed at altering growth rates may be one way to demonstrate this potential fitness tradeoff. In yeast cells exposed to oxidative stress agents, partial deletions within mtDNAs may initially be under genetic control, as a way to quickly reduce endogenous ROS levels by preventing OXPHOS [52]. We found that growth rates and *petite* frequencies were increased by altering environmental conditions. It is possible the increased OXPHOS requirements of rapidly dividing cells may also stimulate this retrograde signaling activity, resulting in higher rates of mtDNA deletions.

In constant environments, there should be rapid selection for variants providing optimal growth rates. This selection may occur more often in yeast where cells are likely to experience singular environments over a generation time. There is some evidence for this. When grown in media attempting to emulate natural habitats, we previously showed that coadapted mitonuclear genome combinations had higher growth rates [15]. Because there is a wide landscape of potential alleles involved in growth, different coadapted mitonuclear loci may evolve rapidly in different populations. In yeast, this could be important in maintaining allelic variation across the species. It will be interesting to see whether growth is a general feature of mitonuclear loci contributing to phenotypes other than mtDNA stability. Given that mitochondria are central to maintaining cellular homeostasis, this may be likely.

#### Main effects

Loci with main effects for mtDNA stability detected by our model interact with mtDNA through non-specific binding. In the MNRC, missense alleles of the mtDNA polymerase, *MIP1*, had the strongest effects on *petite* frequencies. It is well documented that the *MIP1* allele in the yeast reference strain, and other experimentally derived variants, lead to a decrease in mtDNA stability [51, 90, 91]. Our work suggests that naturally occurring variants contribute to basal differences in mtDNA stability in populations. Low expression of the mammalian homolog, PolG, leads to the accumulation of mtDNA deletions [92, 93]. Consistent with this, we found that expression of *MIP1* was lower in strains with the low *petite* frequency alleles and positively correlated with *petite* frequencies (**Fig S6, S7**), although not to statistical significance. Interestingly, *MIP1* expression was dependent on the genotypes of the mitonuclear associated genes, and *BNS1* expression was dependent on the genotype of *MIP1* (**Table S12**). From this data, it can’t be deduced whether cells alter *MIP1* expression in response to growth differences caused by genotypes of mitonuclear loci or if expression of mitonuclear genes is regulated by Mip1 levels. Significant mitonuclear genotype × *MIP1* genotype interactions influenced the expression of the mitonuclear associated genes *BNS1* and *YAT1* (**Table S12**) and suggest that higher order interactions are involved in mtDNA stabilities.

We also found that *EST1* variants associated with mtDNA stability independently of mitotype and that its deletion altered *petite* frequencies. Est1 maintains telomerase at the linear ends of nuclear chromosomes [94, 95] by stabilizing G-quadruplex secondary structures in the DNA [96]. G-quadruplexes play a role in stabilizing human mtDNAs and can cause mtDNA polymerase to stall [97]. The GC-clusters in yeast mtDNAs likely form secondary structures [63] though it is not known whether these form typical G-quad structures or whether they act to stabilize mtDNAs. While Est1 is normally targeted to the nucleus, a proteomics experiment found that Est1 coimmunoprecipitated with mitochondrial ribosomes just prior to cell division (in G2 phase) [98]. It is plausible that Est1 plays dual roles in nuclear and mitochondrial genome maintenance. Another non-specific mtDNA binding protein identified by our association model (but not verified here) was *ILV5*, a mitochondrial nucleoid associated protein that was previously shown to influence mtDNA stability [99].

While higher GC content typically stabilizes DNAs, we found that mtDNAs with the highest GC-content were the least stable (**Fig 2A**). This is consistent with the observation that *petite* frequency differences between different yeast species correlates with their GC-cluster content [100]. Within *S. cerevisiae*, the M4 family of GC-clusters appears to be expanding in the mtDNAs with West African lineages by targeting another family of clusters [67] and likely explains the high *petite* frequencies in these strains (**Fig 2B**). Not much is known about functional differences between different categories of these mobile elements [63, 66]. Previously, we carefully aligned 9 mtDNAs, including their long intergenic regions, and showed that only a small number of GC clusters were in conserved positions [67]. Interestingly, the M4 clusters in the West African mtDNA in that alignment interrupted 15% (2 of 13) of the conserved clusters. The M4 clusters could destabilize mtDNAs directly or perhaps their expansion has interrupted genome stabilizing functions provided by the positions of the conserved clusters.

### Significance

By focusing on naturally occurring genetic variation in *S. cerevisiae*, the alleles mapped using the MNRC provide insight into evolutionary potential. We found that mitonuclear interactions affecting mitotic growth contribute to mtDNA stability although the significance of these specific interactions in yeast populations is not yet known and will depend on allele frequencies and relative effect sizes in context with environments in mating populations. Mitochondrial and nuclear sequences follow different evolutionary trajectories in yeast populations [101] and much of the genetic variation is unique to a single population [47]. It is likely that the MNRC contains combinations of interacting loci that do not exist in nature. Large scale intra- and interpopulation surveys will be required to determine how selection on mitonuclear interactions has shaped patterns of diversity across geographic ranges and ecological niches. Thus, it is not clear that the alleles identified here contribute to population specific phenotypic differences but it is clear that mitonuclear interactions have shaped phenotypic diversity. In addition, the mitonuclear loci identified using the MNRC provide insight into pathways and mechanisms that generate phenotypic differences. The MNRC offers a powerful tool to identify mitonuclear interactions and helps us better understand and predict the complex genotype/phenotype relationships that shape life.

## Methods

### Yeast strains

All strains are listed in **Table S1**. Wild yeast isolates, described in [102], were obtained from the National Collection of Yeast Cultures General. Deletion strains [103] were obtained from the Yeast Knockout Collection (Horizon Discovery). To replace mtDNAs, karyogamy deficient matings were performed as previously described [15].

### Media

Media recipes include: SD (6.7 g/L yeast nitrogen base without amino acids, 20 g/L glucose); CSM with or without amino acids as specified (SD + 800mg/L CSM premix or as recommended by the manufacturer (Sunrise Science)); (CSM containing 30mL/L ethanol and 30mL/L glycerol instead of glucose); sporulation media (1% IOAc, 0,1% yeast extract, 0.05% dextrose); YPD (10g/L yeast extract, 20g/L peptone, 20g/L glucose) supplemented with 10 mM CuSO4 when indicated; YPEG (YPD + 30mL/L ethanol, 30mL/L glycerol instead of glucose); YPDG (1% yeast extract, 2% peptone, 0.1% glucose, 3% glycerol). Sugar concentrations in CSM media were altered as indicated. For solid media, agar was added to 2% prior to autoclaving.

### Petite *frequency assays*

To assay rates of mtDNA deletions, 5 mL YPD cultures were inoculated with freshly grown colonies, grown in roller tubes at 30°C for exactly 15.0 hrs, diluted and plated onto YPDG solid media. After 2-3 days at 30°C, large (*grande*) and small (*petite*) colonies were counted manually or photographed and counted using an imaging system (sp-Imager-SA, S&P Robotics, Inc). To quantify *petite* frequencies in the deletion strains, 20 freshly grown colonies were scooped from solid media, diluted and plated directly onto YPDG solid media. Single assays were performed for each strain in the recombinant collections. All other assays were performed in 3-20 replicates. *Petite* frequencies are reported as the ratio of *petite* to *grande* ([#*petite colonies/ total colonies*] *\* 100*).

### Growth phenotyping

To test to the effects of sugar and temperature on growth, strains were cultivated in 96 well microtiter plates using Biotek Eon photospectrometers using double orbital shaking. Optical densities (600nm) were recorded at 15-minute intervals until cells reached stationary phase. Maximal growth rates (V_max_) were determined as the highest slope of regression lines modeled over sliding windows of 5 data points from growth curves and normalized to the mean of a reference strain included on each plate.

### Multiparent recombinant strain collection

An F7 multiparent recombinant collection of *S. cerevisiae* yeast derived from 25 wild isolates and paired with 3 different mtDNAs was created for genome-wide association studies. We first created a set of parental strains containing a single mitotype with selectable markers to facilitate matings. We began with haploid derivatives (*MAT*a and *MATα ura3∷KanMX)* representing 25 wild yeast isolates (**Table S1**). To facilitate the selection of diploids between these haploid strains, an *arg8∷URA3* disruption cassette from *BamHI* linearized plasmid, pSS1, was introduced into each *MATα* strain through chemical competence (EZ Yeast Transformation Kit (Zymo Research) or [104]) or electroporation [105]. Transformants (strains MLx2×1UA-MLx28×1UA) were selected on CSM-ura solid media and arginine auxotrophies and respiratory competencies were verified by replica plating to CMS-arg and YPEG, respectively. Correct integration of the *arg8∷URA3* disruption cassette was verified through tetrad analysis: each *MATα arg8∷URA3* strain was mated to their *MATa ura3∷KanMX* isogenic counterpart, diploids were selected on SD media and sporulated on SPO media. Spores were dissected from ≥ 12 tetrads and printed to CSM-ura, CSM-arg, and YPEG media, verifying a 2:2 segregation of Arg− Ura+: Arg+ Ura− phenotypes. All crosses had >90% spore viabilities. The mtDNAs in the *MAT*a strains were replaced with the mtDNA from 273614N using karyogamy-deficient matings as previously described [15]. The mtDNAs from the *MATα arg8∷URA3* strains were removed through ethidium bromide treatment.

To create the multiparent recombinant collection, the MATa *ura3∷KanMX* ρ^+^ and MAT*α ura3∷KanMX arg8∷URA3* ρ^0^ strains were mated to create each possible heterozygous F1 diploid. To do this, aliquots (50 μL) of haploid strains were mixed with 200 μL fresh YPD media and incubated without mixing for 2 days at 30°C in 96-well plates. Mating mixtures were harvested by centrifugation, washed (200 μL ddH_2_0), re-suspended in 250 μL CSM-URA-ARG liquid media and incubated for 2 days at 30°C with continuous shaking, and then spotted onto solid CSM-URA-ARG media. This yielded 279 F1 heterozygous diploids (of 284 attempted crosses). To create F1 haploids, 10 μL aliquots of diploid-enriched mixtures were spotted to sporulation media and incubated at room temperature until tetrads were visible via compound microscopy (7-11 days). The sporulated cells were collected and pooled by washing the plates with 5mL ddH_2_0. To isolate spores from asci and vegetative cells, a random spore analysis protocol was followed [106], with modifications. Cell walls were digested by incubating 1 mL aliquots of the sporulated cells with zymolyase 20T (1 mg/ml)) at room temperature for 1 hr with gentle rocking. The cells were centrifuged (12000 rpm, 4°C, 1min), washed (1 mL ddH_2_0) and resuspended in 100 μL ddH_2_0. The treated cell mixture was vortexed vigorously for 3 minutes to adhere spores to tube walls. The supernatant was carefully aspirated, and the spores were gently washed (1mL ddH_2_0) to remove remaining vegetative cells. To release spores from the plastic tube walls, cells were sonicated for 10-20 sec at 110 V (Stamina XP s50.0.7L, SharperTek) in 1mL of 0.02% Triton-X. The released spores were centrifuged (12000 rpm, 4°C, 1min), washed (1mL ddH_2_0), and re-pelleted. The resulting freed spore mixtures were combined into a single tube with 1mL ddH_2_0. Spore density was determined with a hemocytomer and plated for single F1 haploid colonies (~500 CFU/plate) onto 10 petri plates containing CSM-URA media (selecting for *MATa* or *α ura3 arg8∷URA3*) and 10 petri plates containing CMS-ARG media (selecting for *MATa* or *α ura3 ARG8*). The F1 haploid cells were pooled by washing the colonies from each plate using ~5 mL ddH_2_0. The pooled cells were washed in 5 mL ddH_2_0, resuspended in 5mL YPD media and incubated at 30°C without shaking. The random mating mixtures were washed in 5 mL ddH_2_0, resuspended in 5 mL CSM-URA-ARG and incubated for 8-12 hours to enrich for F2 diploids. The cell mixtures were pelleted, resuspended in 2.5 mL ddH_2_0, aliquoted (250 ul) onto solid sporulation media and incubated at room temperature for 7 days. Spores were released and F2 haploids were isolated as described above. Random matings and haploid selection were repeated for a total of 7 meioses. Haploid F7 progeny were isolated as single colonies on YPD media, and replica plated to CSM-ARG, CSM-URA, and YPEG to determine auxotrophies and respiratory growth. Mating types were tested using mating type testers. Approximately 200 verified haploid recombinants were selected as strains for Recombinant Collection 1 (RC1) and include ~50 isolates of each selectable genotype. RC001-RC049: *MATa ura3 ARG8;* RC102-146: MATα *ura3 arg8∷URA3;* RC201-247: *MAT*a *ura3 arg8∷URA3;* RC301-347: MATα *ura3 ARG8*.

The mtDNAs from each strain in RC1 were removed using an ethidium bromide treatment (creating RC001-347 *ρ*^0^) and replaced using karyogamy deficient matings to create RC2 and RC3. RC2 strains contain the mtDNA from YPS606 and were created using mitochondrial donor strains SPK27 (for RC2:001-048), CK520E1 (for RC2:102-146 and RC2:301-347) and MKG109 (for RC2:301-307). RC3 strains contain the mtDNA from NCYC110 and were created using mtDNA donor strain TUC131.

### Genome Sequencing and analysis

Genomic DNAs from 192 F7 recombinants from RC1 were isolated following Hoffman-Winston genomic DNA protocol (Hoffman & Winston 1987). DNA samples were concentrated using ZR-96 Genomic DNA Clean & Concentrator™ – 5 (Zymo Research Corp). DNA concentrations were measured using Qubit and diluted to the final concentration of 0.2 ng/μl. DNA sequencing libraries were prepared using an Illumina Nextera XT DNA Library Prep Kit according to manufacture instructions. The DNA concentration of each library was determined using Qubit, normalized, and pooled. The pooled libraries were sequenced using a single run of paired end 2 × 150 bp on Nexseq500 (Illumina) at the Institute for Biotechnology and Life Sciences Technology at Cornell University. The reads for each RC strain were individually mapped to the annotated reference genome S288C (R64-2-1_20110203) using Bowtie2. Single nucleotide variants (SNPs) and regions of low and high coverage were identified using the Find Variations/SNPs function in Geneious v.8, filtering the results to regions with a minimum coverage ≥5, minimum variant frequencies within the reads ≥ 0.8 and the maximum variant *P* values (the probability of a sequencing error) ≤10^−6^. To create a SNP table for association mapping, adjacent polymorphisms were merged, telomeric regions were removed and polymorphisms from each strain were combined into a single file. Low coverage areas were converted to deletions and the polymorphisms were filtered for biallelic variants with MAF > 0.5%. This resulted in in 24,955 polymorphic sites.

The mtDNA sequences were analyzed for GC cluster and intron content as previously described [67]. Accession numbers are provided in **Table S4**.

### Statistical analysis and association testing

Statistics and plotting were performed using R 4.1.2 [107] and association models were performed using R 4.0.4 in RStudio [108]. Generalized linear models of the family binomial accounting for the different numbers of *petite* and *grande* colonies were performed using *glm* with the *cbind* function. ANOVAs for the analyses of growth or expression were performed using linear models with the lm function. Correlations were run using the *corr.test* function. Gene ontology classes for associated genes were identified using GO∷TermFinder (Boyle, 2004 https://doi.org/10.1093/bioinformatics/bth456) and tested for over-representation using Fisher’s exact tests.

### Genome-wide association analysis

A biallelic SNP table containing 24955 unique SNPs was used for association testing. To address residual population structure in the recombinant strains, a principal component analysis was performed using PLINK v1.9 [109]. A simple association test was performed on the growth parameters from RC1 strains grown on YPD + 10 mM CuSO4 using the linear model: *colony size ~ covariates + nuclear SNP + error* where colony size is the strain mean derived from the linear model, nuclear SNP represented SNP variation at a given locus, and covariates included auxotrophies, mating type, and the first ten principal components from the PCA analysis. Association tests to detect mitonuclear interactions were performed using the strain means for *petite* frequency from RC1, RC2 and RC3 in the following model∷ *phenotype ~ covariates + nuclear SNP + mtDNA + nuclear SNP*mtDNA+ error*, where phenotype is either the petite frequencies. Covariates included MIP1 alleles when indicated. False discovery rates were calculated using the “*qvalue”* package version 2.22.0 [110].

Associated SNPs that crossed FDR threshold were further filtered to non-synonymous SNPs within coding sequence (CDS) regions, and upstream regions within 250bp upstream of the CDS. Effect sizes for the associated SNPs with main effects were determined as the absolute differences in *petite* frequencies for strains with each variant weighted by their allele frequencies (effect size =Δ= |(freq _allele1_ × *petite* freq_allele1_) – (f _allele2_ × *petite* freq_allele2_)|).

### RT-qPCR

Cells from overnight CSM cultures were lysed according to [111] and total RNA was isolated using Qiagen RNeasy mini spin columns. RNA was quantified (Invitrogen Qubit Fluorometer) and diluted to 92 ng/μL. BioRad iScript cDNA Synthesis Kit was used to create cDNA, according to manufacturer’s instructions. RT-qPCR was performed on a BioRad CFX Connect Real-Time PCR Detection System by adding 2 μl (~2.3 ng/μl) cDNA to 10 ul of BioRad SsoAdvanced Univerisal SYBR Green Supermix, 1 μL each of forward and reverse primers (500 nM) and 6 uL nuclease-free H_2_0 in BioRad Hard-Shell 96-Well PCR Plates according to the following protocol: [95C-30s, (95C-15s, 60C 15-40s)_40_, 65C-95C-0.5C/5s]. Primer sequences are provided in **Table S14**. Relative expression was determined as the residuals from a linear regression of 1/Ct for each candidate gene against 1/Ct for the control gene, UBC6 as previously described [112]. This approach controls for differences in RNA extraction and cDNA manufacturing efficiencies and results in more normally distributed data where higher residual values correspond to higher starting mRNA levels.

## Data Availability

Reads for the MNRC strains can be retrieved from NCBI under the BioProject ID PRJNA871925. Scripts, SNP table and data available at https://github.com/mito32/Mitonuclear-Recombinant-Collection.

## Author Contributions

HLF, ACF, and KC conceived the project. ML, MG, and FR created the recombinant collection. Library preps were performed by THMN, JFW, and WL. Phenotyping was performed by THMN, MG, FR, WL, ATB, MS, BB, MT, BG. Data analysis was performed by THMN, ATB, JFW, ACF and HLF. The manuscript is written by THMN, ACF and HLF. All authors read and improved the manuscript.

## Acknowledgements

We gratefully acknowledge T.D. Fox for the gift of plasmid, pSS1 and L. P. Musselman for assistance with RT-qPCR. We are also grateful to Binghamton University undergraduate students K. Dave, B. DeJesus, J. DeJesus, A. Federico, G. Lal, S. Sondhi, S. Oh, B.A. Wong, and A. Ziesel for technical assistance with *petite* frequencies in RC1.

## Supplemental Tables

**Table S1.** Strain table

**Table S2.** ANOVA *Petite* frequencies in wild isolates vary by strain and population

**Table S3.** ANOVA mtDNAs influence *petite* frequencies in the Y55 nuclear background

**Table S4.** mtDNA GC clusters and introns

**Table S5.** ANOVA *Petite* frequencies influenced by mitonuclear interactions in a 4×4 mitonuclear genotype panel

**Table S6.** Recombinant Collection SNP descriptions

**Table S7.** Associated SNPs that influence *petite* frequencies independent of mitotype (main effect loci)

**Table S8.** Associated SNPs that influence *petite* frequencies dependent on mitotype (mitonuclear loci)

**Table S9.** Effect sizes of main effect loci

**Table S10.** Effect sizes of mitonuclear loci

**Table S11.** ANOVAs Mitonuclear effects of *HGH1* and *YAT1*

**Table S12.** ANOVA Mitonuclear effects of *HGH1*, *BSN1*, *SMI1*

**Table S13.** ANOVAs Expression differences of mitonuclear candidate loci

**Table S14.** Primer sequences for qRT-PCR

## Supplemental Figures

**Fig S1.**
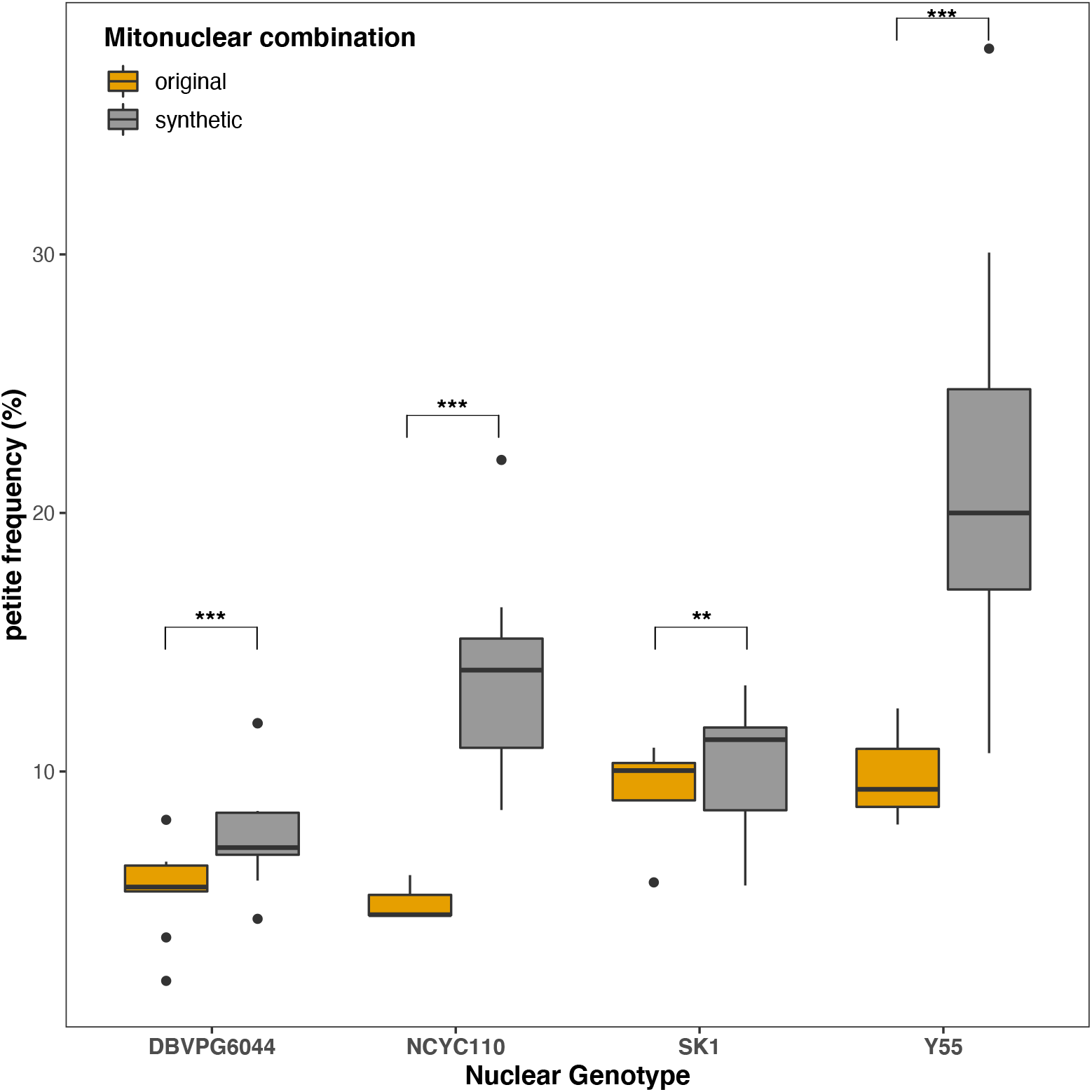
Strains harboring original mtDNAs have lower *petite* frequencies than those with synthetic mitonuclear combinations. *Petite* frequencies of strains with the original (gold) vs. synthetic (grey) mitonuclear genotypes from **Fig 3** are replotted as box plots with the synthetic combinations combined. All nuclear and mtDNAs are from strains with West African lineages. ANOVA significances are shown. * *P<0.05*, ** *P* ≤ 0.005, *** *P* ≤ 0.001.

**Fig S2.**
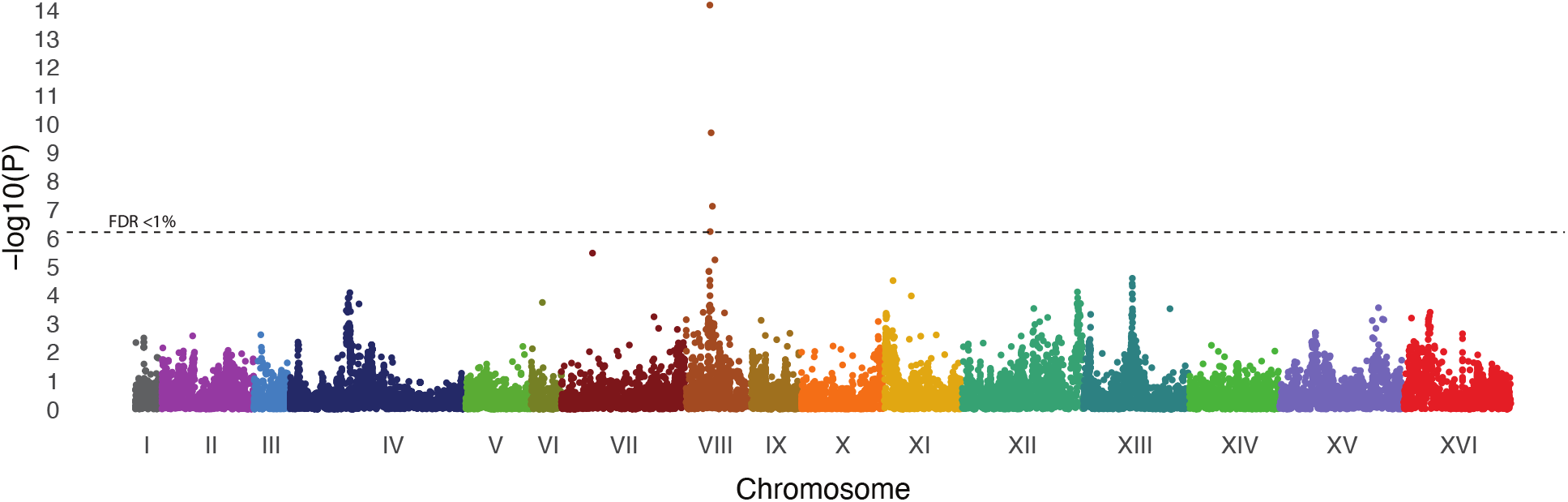
Association Mapping using the Recombinant Collection. A Manhattan plot of −log10 of *P* values plotted against chromosomal position shows associations for maximal colony sizes for RC1 strains grown on copper sulfate. A single peak, corresponding to the location of *CUP1* on Chr. 8, is the only significant association at FDR<1%.

**Fig S3.**
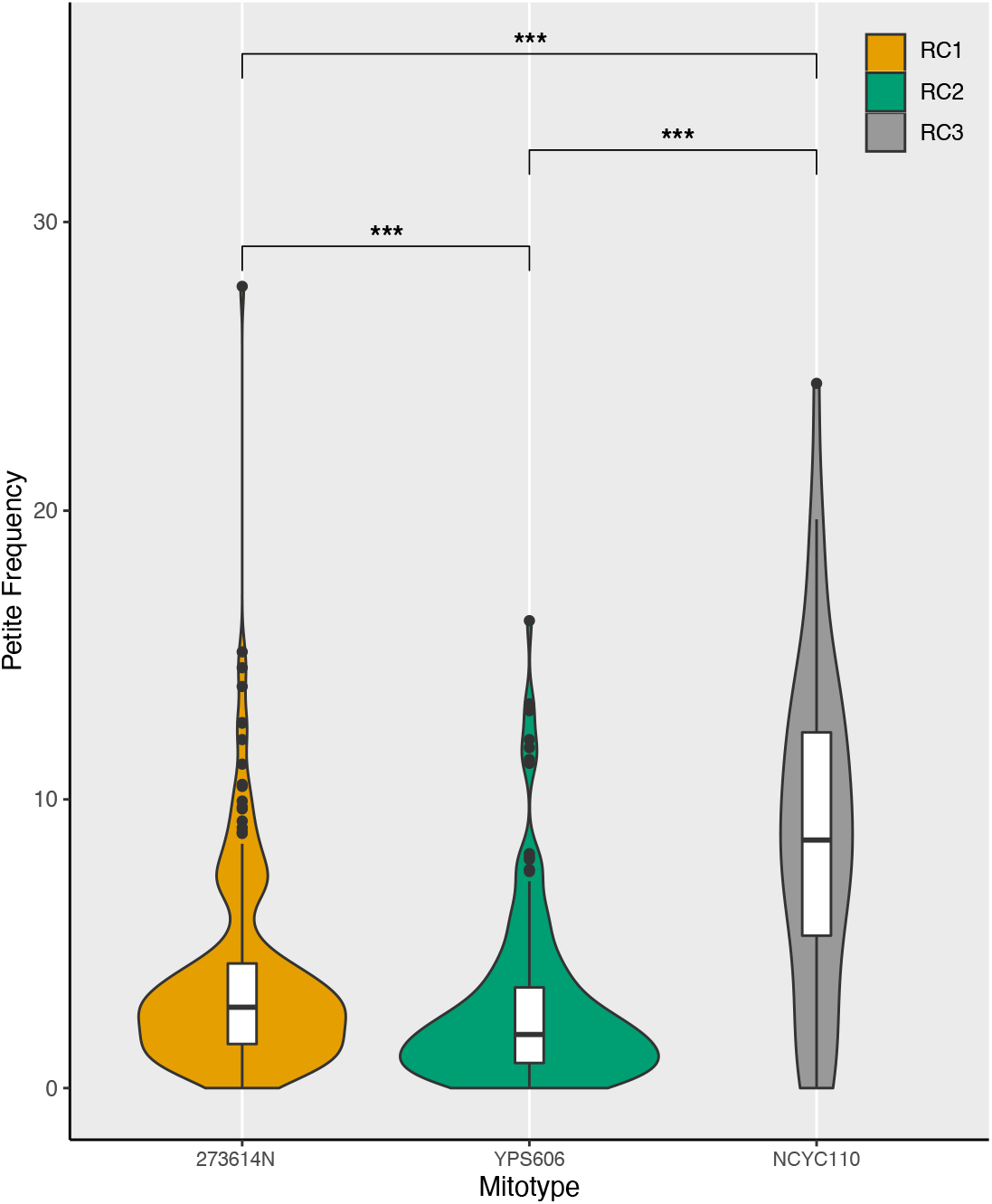
Strains with the GC-rich mtDNA in RC3 strains have higher *petite* frequencies. *Petite* frequencies of strains from RC1, RC2, and RC3 are presented as violin plots. RC1 strains harbor mtDNA from 273614N containing a medium number (137) of GC clusters. RC2 strains harbor mtDNA from YPS606 containing a low number (117) of GC clusters. RC3 harbor mtDNA from NCYC110 containing a high number (210) of GC clusters.

**Fig S4.**
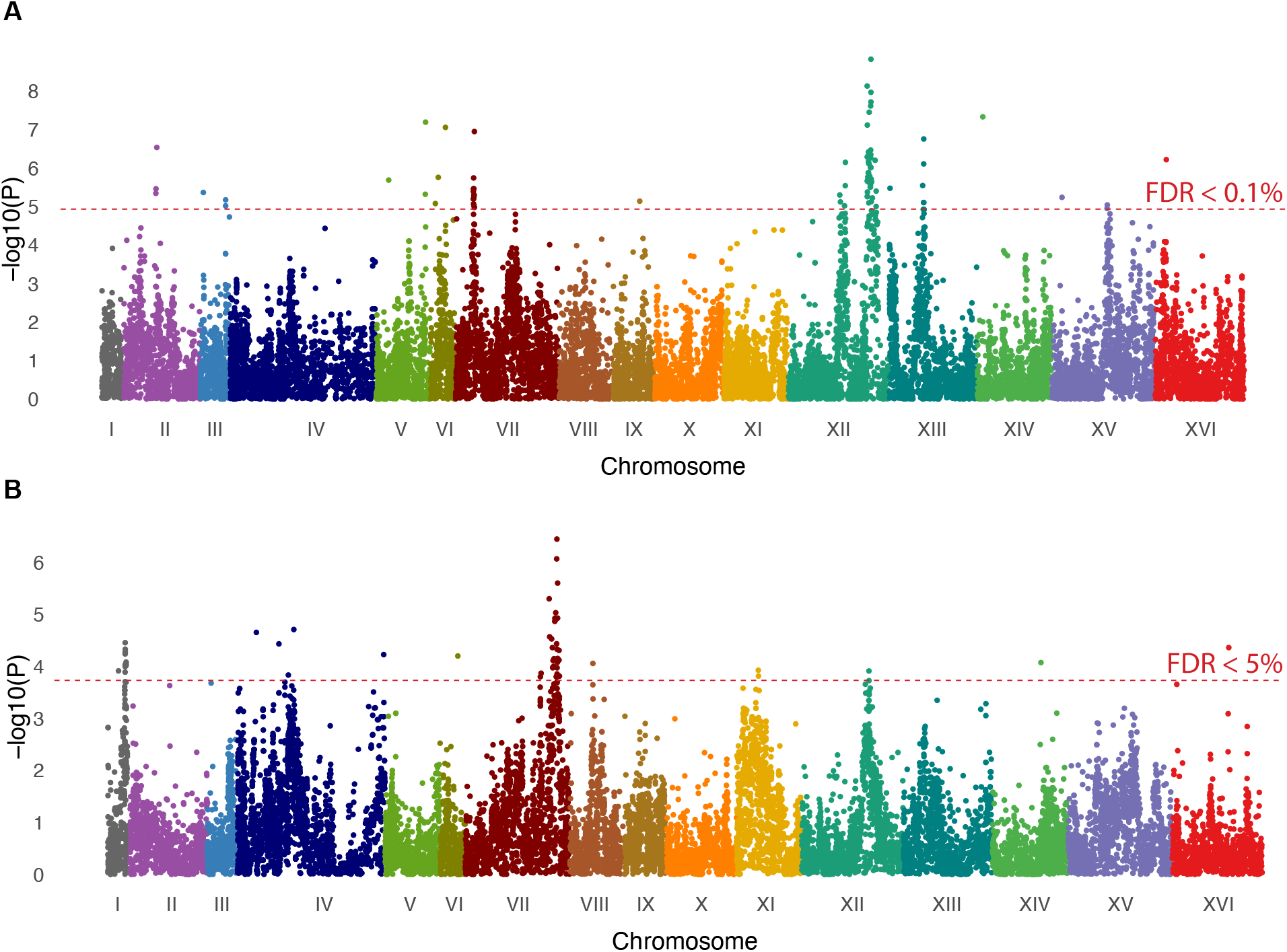
GWAS using MIP1 variants as covariates increases power to detect mitonuclear associations. Manhattan plots show mitotype **A**. independent and **B**. mitotype dependent associations, when *MIP1* variants were included as covariates. Red lines indicate FDR thresholds at 0.1% for nuclear associations and 5.0% for mitonuclear associations.

**Fig S5.**
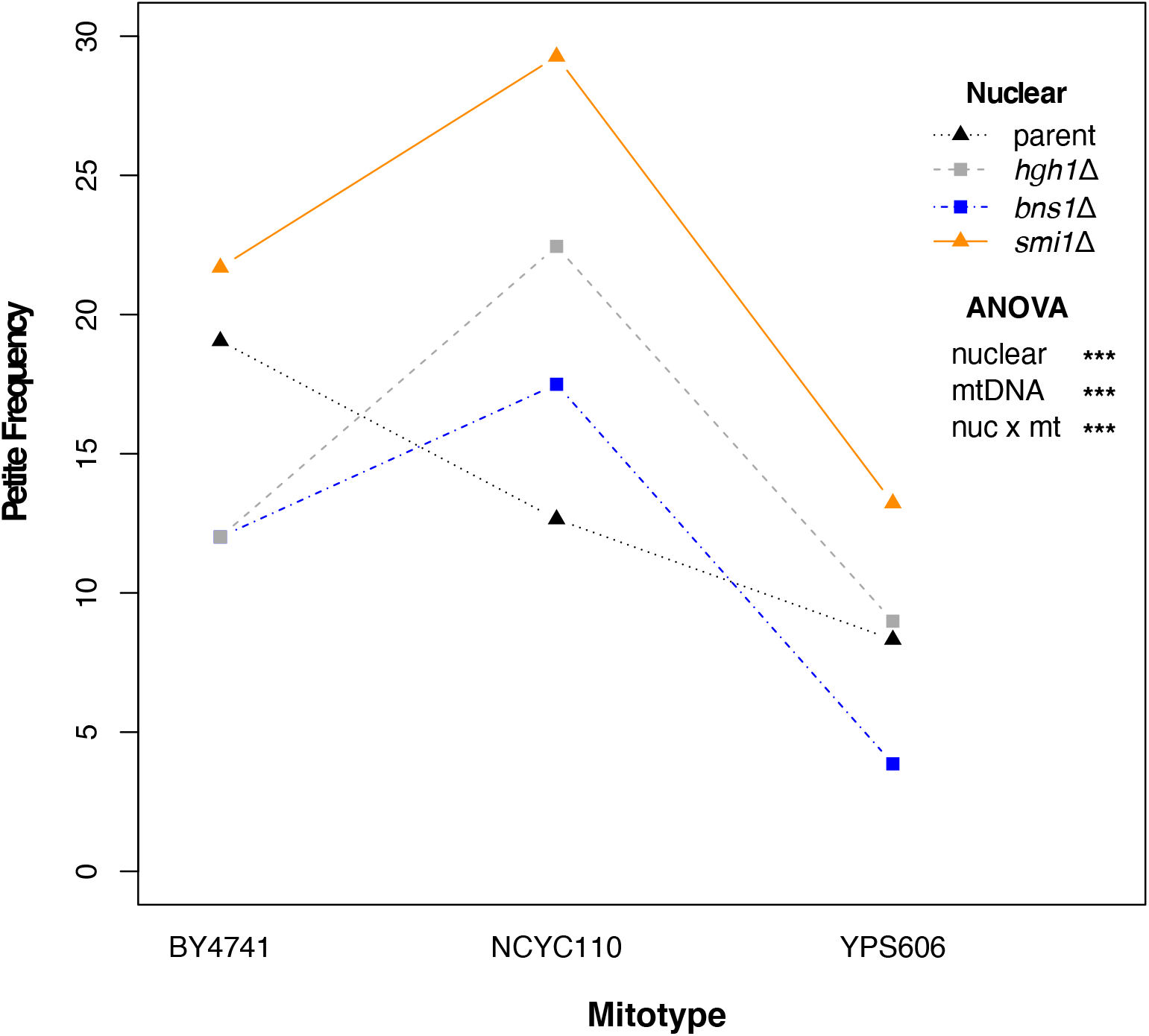
Mitonuclear interactions of BNS1, SMI1 and HGH1 on *petite* frequencies. Interaction plot follows *petite* frequencies for each nuclear genotype paired with different mtDNAs. See **Table S11** for ANOVA. Data was collected using the same assay as performed for phenotyping the RCs (with 8-12 replicates) and cannot be combined with the data shown in **Fig 6**.

**Fig S6.**
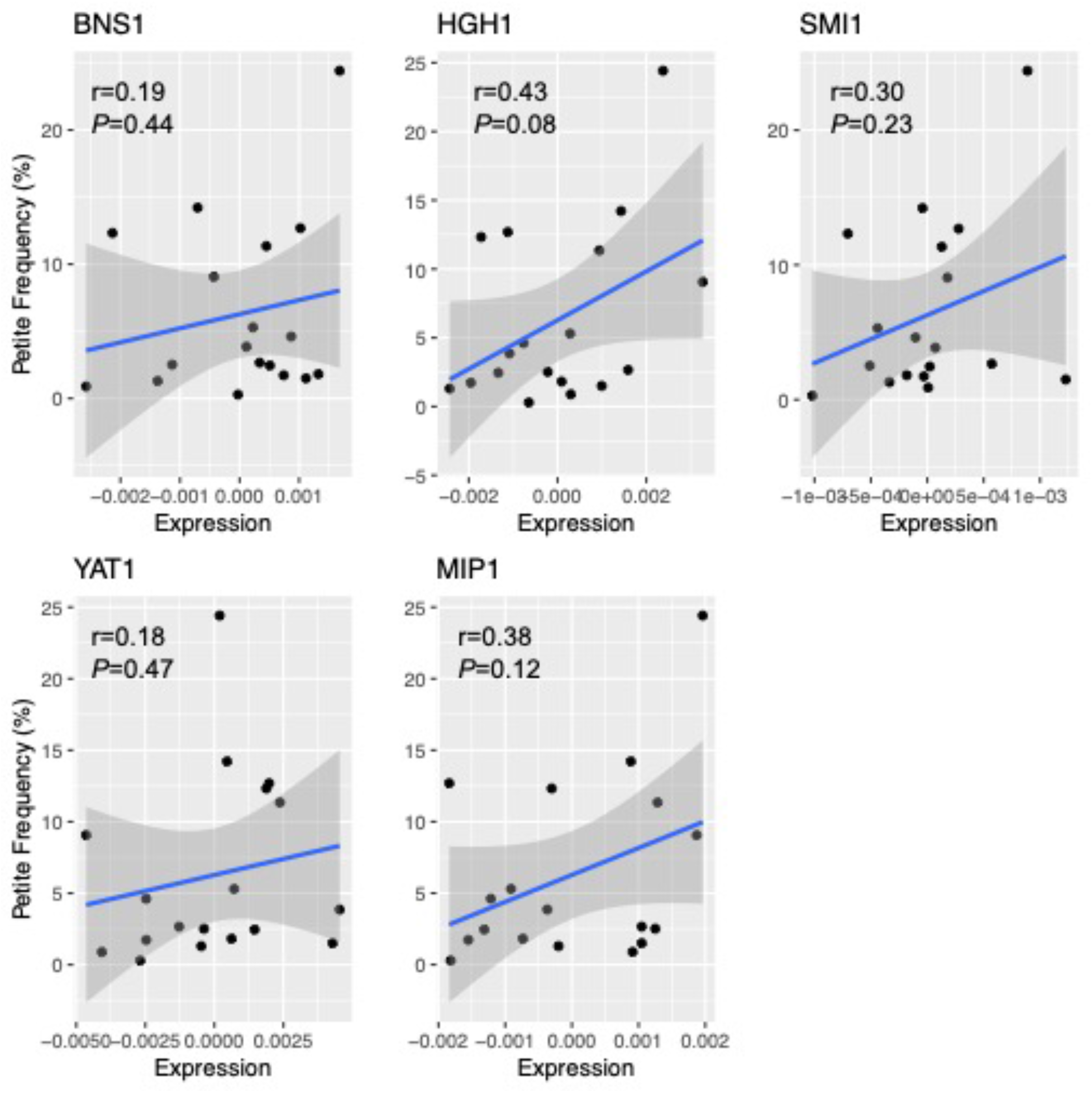
*Petite* frequencies do not correlate with mRNA expression of associated genes. Normalized expression levels (as residuals from regression lines of mRNA levels of each gene compared to a control gene) were plotted against *petite* frequencies. All genes showed positive correlation with *petite* frequencies, though no correlation was statistically significant.

**Fig S7.**
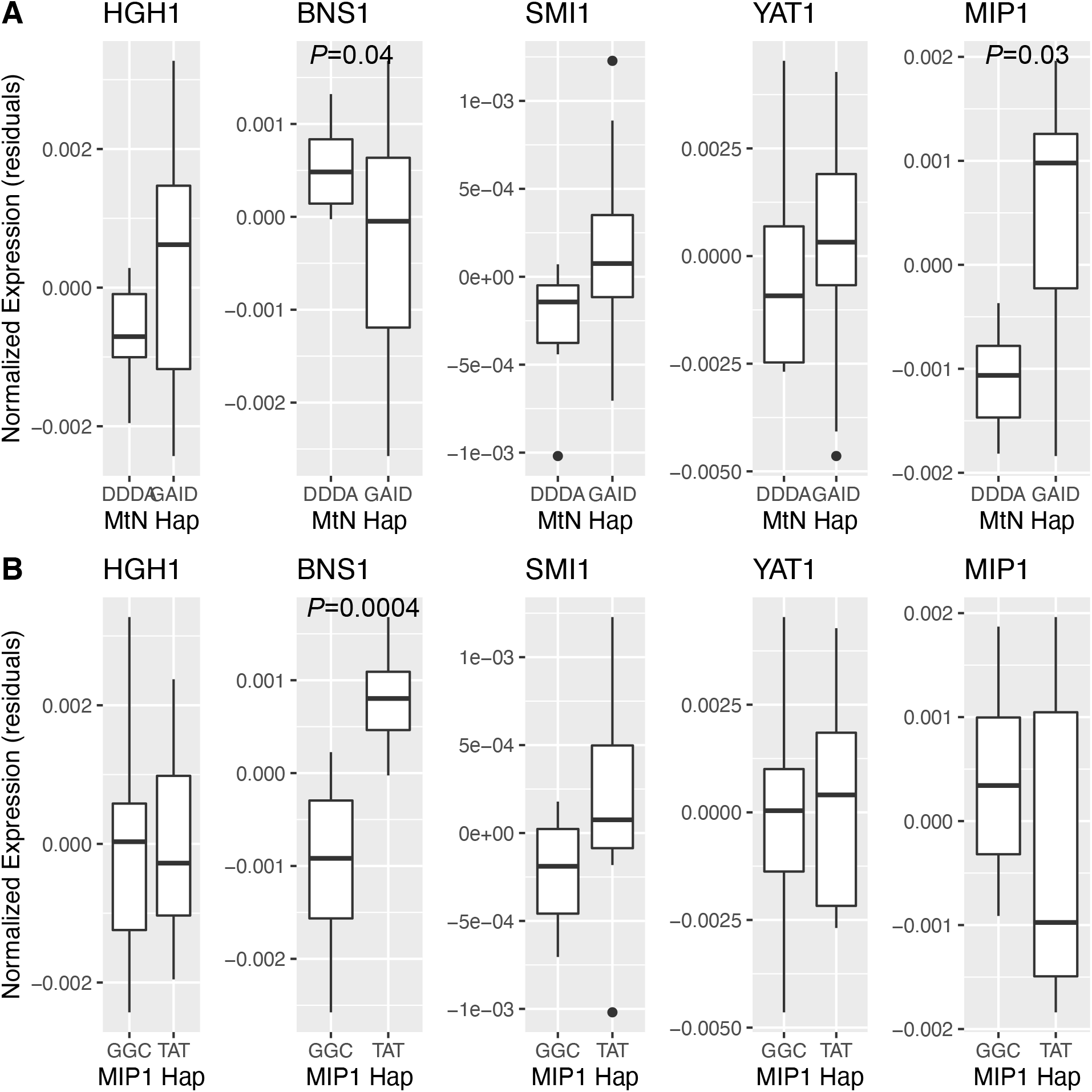
Expression of candidate genes by nuclear haplotype. **A.** Normalized expression levels of each candidate gene separated by haplotypes of **A**. mitonuclear candidate loci or **B**. *MIP1* loci. The mitonuclear haplotypes represent the SNPs with highest effect sizes for each candidate gene. *P* values for significant differences are shown. All other comparisons were non-significant.

**Fig S8.**
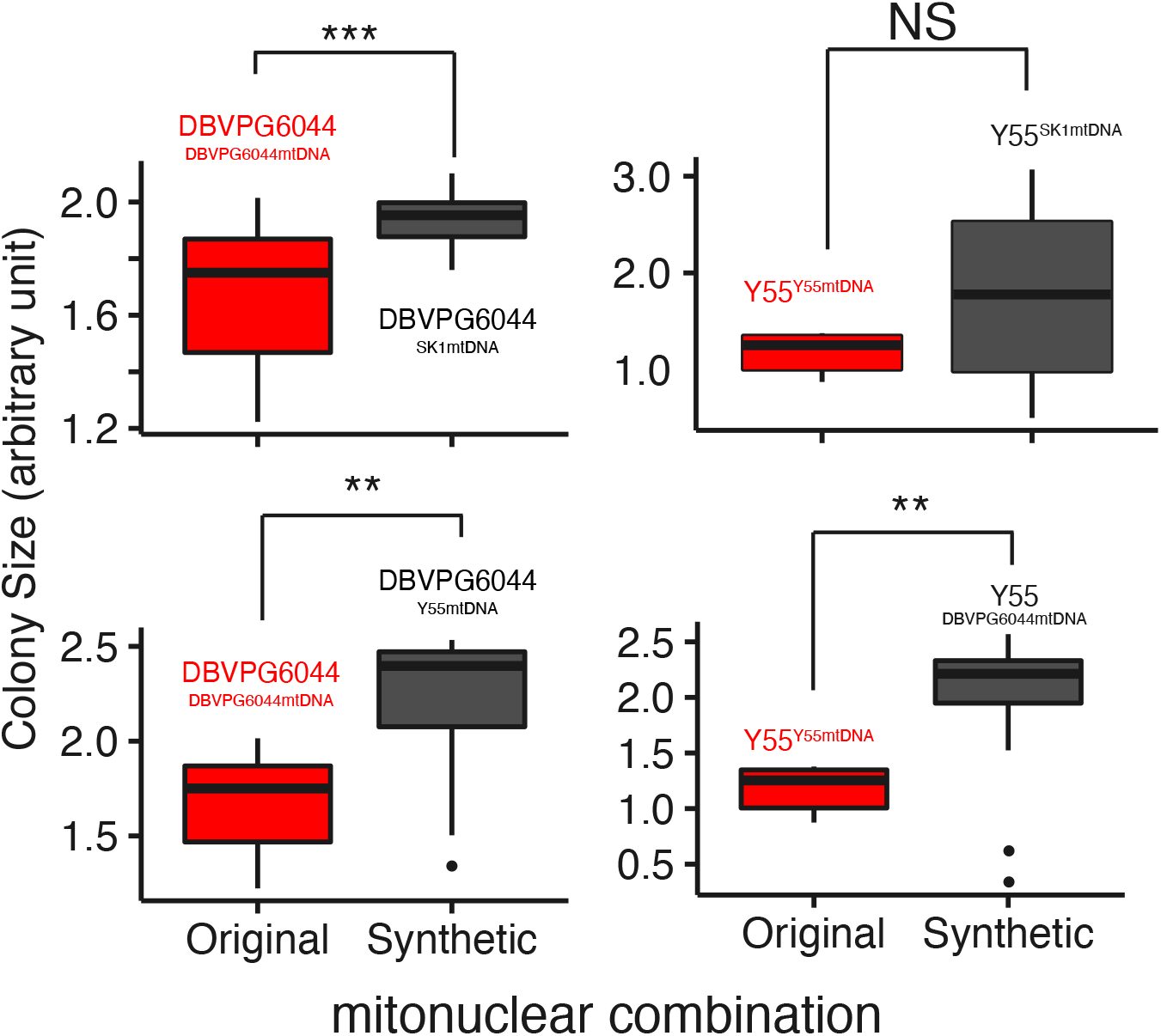
Synthetic mitonuclear genotypes with increased *petite* frequencies have increased growth rates. Maximum colony sizes for strains containing original or synthetic mitonuclear genotypes are presented as boxplots. Each synthetic mitonuclear genotype had higher growth, and higher *petite* frequencies (**Fig S1**), than the original mitonuclear genotype. Growth data were from NGUYEN *et al*. 2020 and collected in the same conditions as the *petite* assays were performed. * *P<0.05*, ** *P* ≤ 0.005, *** *P* ≤ 0.001

**Figure S9.**
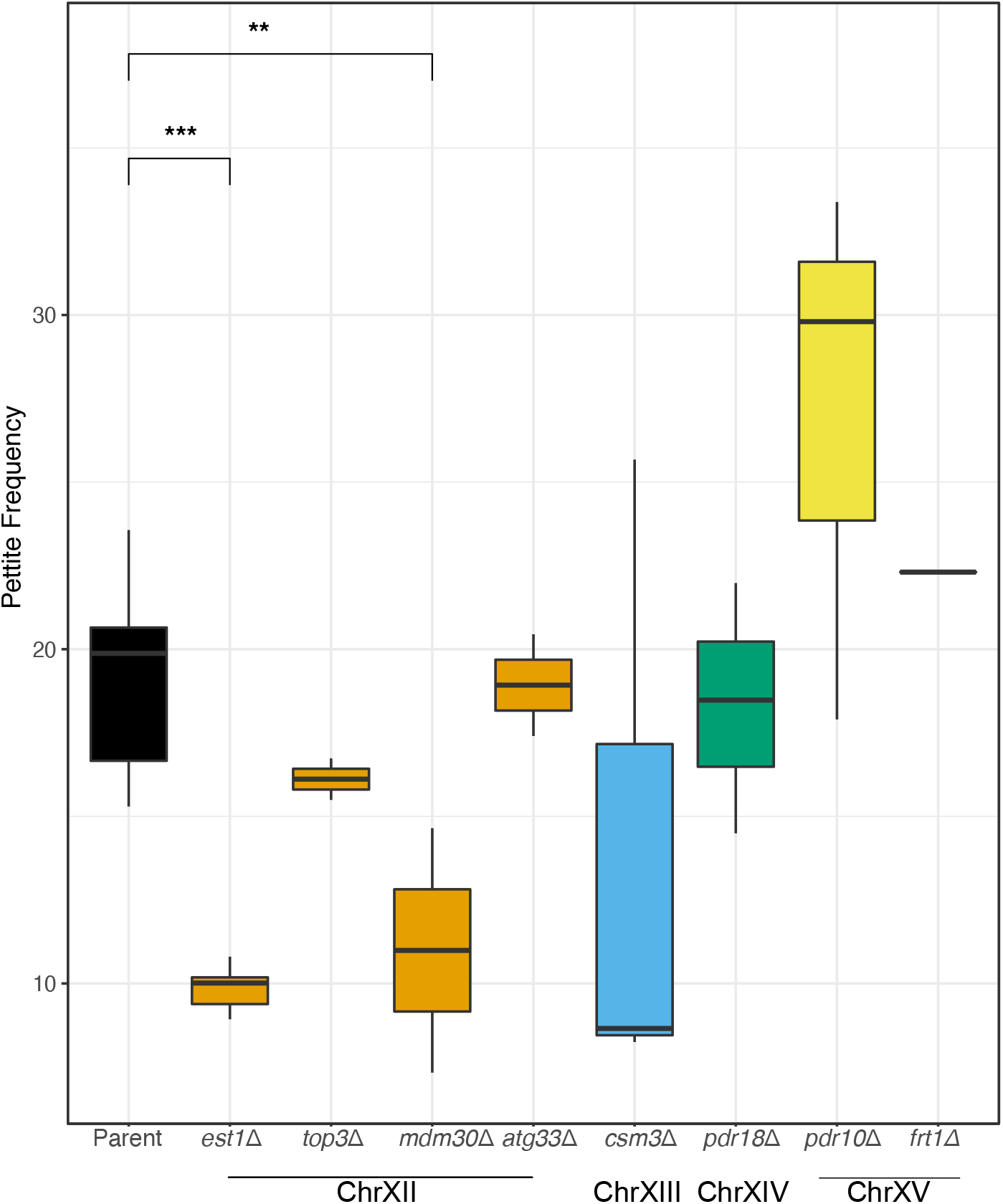
*Petite* frequency of candidate genes associated with mtDNA stability. *Petite* frequencies of strains containing deletions of candidate genes that did not depend on mitotype are shown as boxplots. Significant differences between the *petite* frequencies of the parental strain and each gene disruption, based on 3 replicates for each strain, are shown. Colors indicate chromosomal location of genes. * *P<0.05*, ** *P* ≤ 0.005, *** *P* ≤ 0.001

